# Contrastive Poincaré Maps: uncovering developmental lineages from single-cell data with scalable hyperbolic embeddings

**DOI:** 10.1101/2025.08.22.671789

**Authors:** Nithya Bhasker, Hattie Chung, Louis Boucherie, Vladislav Kim, Stefanie Speidel, Melanie Weber

## Abstract

Embedding hierarchical data in low dimensions is a problem common to many domains: the number of distinct states grows exponentially with depth, whereas Euclidean area grows only polynomially with radius, so branches crowd, cross, and collapse. Hyperbolic geometry, whose area grows exponentially with radius, matches this expansion. Developmental single-cell RNA sequencing (scRNA-seq) data is a demanding instance of this problem, and one in which embeddings are the primary lens on lineage structure, yet the dominant tools, t-SNE, UMAP, and PHATE, are Euclidean. Existing hyperbolic methods such as Poincaré Maps (PM) rely on precomputed pairwise similarity matrices and hand-tuned manifold metrics, scale quadratically in memory, and require PCA preprocessing. We present Contrastive Poincaré Maps (CPM), which learns hyperbolic embeddings directly from raw expression through a contrastive objective, processing data in batches without dimensionality reduction, batch correction, or feature engineering. On a synthetic differentiation tree, CPM uniquely combines faithful global hierarchy with low worst-case distortion, reducing distortion roughly 15-fold relative to PM (1.7 vs. 25.9), while its peak GPU memory stays essentially constant with dataset size and over 120-fold below PM at 25,200 cells. Across two case studies (116,312 mouse gastrulation cells and mouse hematopoiesis), CPM preserves global developmental hierarchies, aligns embedding geometry with biological axes, and supports trajectory analysis without additional processing. By coupling hyperbolic geometry with contrastive learning, CPM delivers scalable, structure-preserving embeddings for developmental scRNA-seq data.

**Code:** https://github.com/NithyaBhasker/ContrastivePoincareMaps

## The Bigger Picture

Branching organization is a recurrent feature of empirical data. A single category subdivides into several categories, each subdivision subdivides in turn, and the population of distinct states multiplies at every successive level. Representing such structures on a flat surface is a question of available area: in a Euclidean (flat) plane, the space accessible at a given distance from the center increases far too slowly to accommodate what recursive branching generates, so that distinct groups overlap and the relations of interest are distorted. This limitation is not an artifact of any particular visualization technique; it follows from the geometry of flat space itself. Hyperbolic space, which carries constant negative curvature, behaves otherwise: the area available expands as one moves outward at approximately the rate branching structures demand. Representations constructed in such a geometry accommodate hierarchies of considerable depth without compressing them.

Existing methods that exploit this property require exhaustive comparison of every pair of items before a representation can be constructed, a requirement that becomes computationally prohibitive once a dataset exceeds approximately one hundred thousand entries. We instead train a model to construct the mapping directly, processing the data in small batches, so that its memory requirement ceases to depend on the size of the dataset. We evaluate the method on cellular differentiation, a domain in which the branching structure has been established independently and therefore furnishes a reference against which any reconstruction may be assessed; the method recovers that structure at a scale at which prior approaches cannot be executed. The model is not itself specific to biology, its input being a table of items and associated measurements. Taxonomies, citation networks, latent structures within document collections, and organizational hierarchies exhibit the same branching organization and encounter the same geometric constraint, and the approach should transfer accordingly.

## 1 Introduction

Hierarchically organized data are common across domains, and rendering them in low dimensions runs into a hard geometric limit. The number of distinct states in a branching hierarchy grows exponentially with depth, whereas the area available in a Euclidean disk grows only as 2*πr*; the space cannot expand fast enough to hold a deep hierarchy, so branches crowd, cross, and collapse, and global relationships are distorted. Hyperbolic space resolves this mismatch directly: its area grows as 2*π* sinh *r*, expanding exponentially with radius and thereby accommodating exponentially many descendants at each level (2–4). A tree embeds into the two-dimensional hyperbolic (Poincaré) disk with low distortion, with radial distance encoding depth and angular position encoding branch identity (Fig. 1).

**Figure 1.**
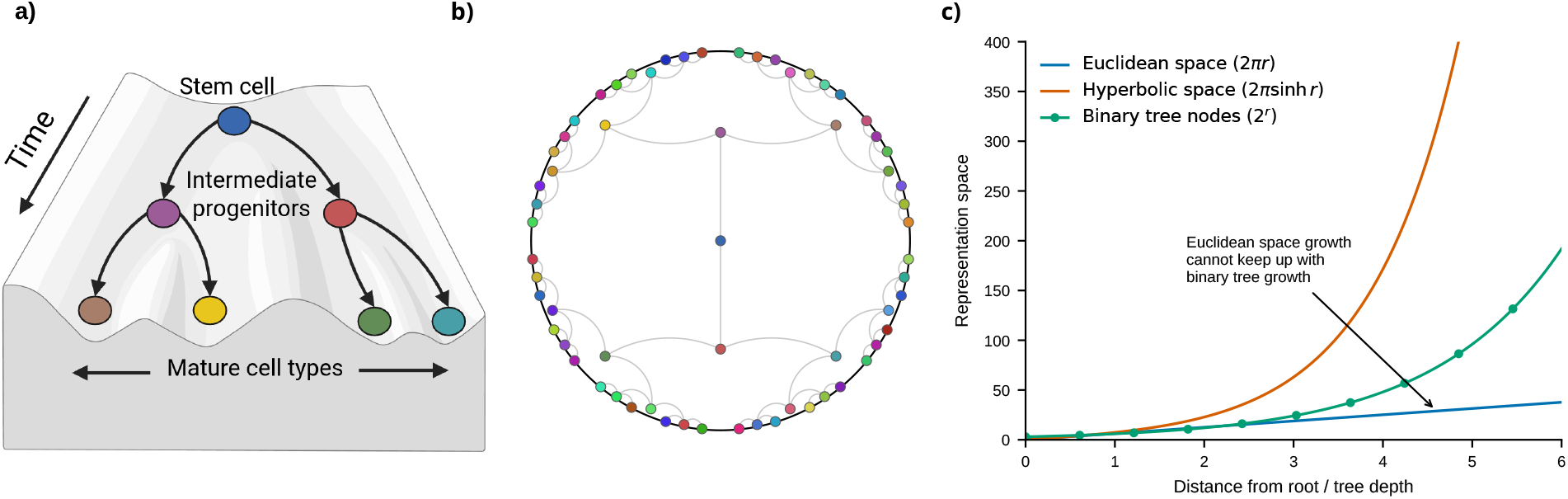
Euclidean space cannot accommodate hierarchical growth; hyperbolic space can. **a)** Waddington’s epigenetic landscape: a pluripotent stem cell differentiates through intermediate progenitors into distinct mature cell types along a branching developmental trajectory. **b)**A branching hierarchy embedded in the Poincaré disk, colored to match panel a: radial distance from the origin encodes developmental depth and angular position encodes branch identity, so successive generations fan out toward the boundary rather than crowding together. **c)**Representation capacity as a function of depth/radius *r*: Euclidean area grows only as 2*πr* (polynomially), while hyperbolic area grows as 2*π* sinh *r* (exponentially), matching the exponential growth of a binary tree (2^*r*^); Euclidean space falls increasingly short of the tree’s growth as *r* increases, while hyperbolic space keeps pace.

Cellular differentiation is a demanding instance of such a hierarchy: pluripotent progenitors give rise to progressively committed states, tracing a branching tree of fates that Waddington famously depicted as descent through an epigenetic landscape. Single-cell RNA sequencing (scRNA-seq) now resolves these lineage trees at the level of individual cells, and low-dimensional embeddings are the primary lens through which biologists read them. Here radial distance becomes developmental depth and angular position becomes branch identity. Yet the dominant tools, t-SNE (5), UMAP (6), and PHATE (7), embed cells in Euclidean space, which creates a central geometric bottleneck in the field.

Hyperbolic embedding of single-cell data is not itself new. Poincaré Maps (PM) established that hierarchies are better preserved in the Poincaré disk than in Euclidean space (8). But PM is a transductive, largely hand-engineered pipeline: it constructs a relative-forest-accessibility similarity matrix over all cells, estimates local and global manifold metrics, and only then opti mizes point positions, behaving effectively as a lookup table that must be recomputed for every dataset. This pairwise construction scales quadratically in the number of cells and exhausts memory on atlas-scale data, and the upstream metric estimation forces PCA preprocessing that discards information and injects its own choices. We remove this machinery. Contrastive Poincaré Maps (CPM) replaces the precomputed similarity matrix and manifold-metric estimation with a learned, contrastive encoder that maps raw expression directly into the Poincaré disk. A projection block performs trainable, end-to-end dimensionality and noise reduction in place of PCA, and a hyperbolic block (9) places cells on the disk. Trained by an InfoNCE contrastive objective (10, 11) with a corruption-based augmentation (12) that pulls together similar cells and pushes apart dissimilar ones, CPM learns robust representations without dimensionality reduction, batch correction, or feature engineering, and processes data in mini-batches so that memory scales linearly rather than quadratically.

This design has two practical consequences. First, because the objective is discriminative rather than generative, CPM optimizes exactly the pairwise structure that downstream lineage analysis depends on, and does not pay the reconstruction cost that generative, ELBO-based embeddings (13) incur when modeling thousands of genes, nor does it suffer the numerical instabilities that prevent the hyperbolic variants of those models from scaling to large datasets. Second, the learned geometry is biologically interpretable by construction: distance from the root measures developmental maturity and angular separation captures differentiation direction. A single CPM embedding therefore supports trajectory analysis, namely pseudotemporal ordering and canonical lineage reconstruction, directly. This replaces a conventional multi-step pipeline, in which highly-variable-gene selection, PCA, batch correction, clustering, neighbor-graph construction, and pseudotime inference each carry their own hyperparameters, with one architecture and one round of tuning.

We validate CPM across synthetic and real developmental data. On synthetic trees with known ground truth, CPM combines faithful global hierarchy with low worst-case distortion and near-constant memory, a balance that competing methods, evaluated with standard neighborhood- and distortion-based metrics (14), fail to achieve: each preserves local rank order, global structure, or distortion at the expense of the others. On mouse hematopoiesis (15), CPM separates the megakaryocyte-erythrocyte and granulocyte-monocyte progenitor lineages and yields root distances that agree with diffusion pseudotime (16) (Spearman *ρ* = 0.75). CPM embeds all 116,312 cells of a mouse gastrulation atlas (17), a scale at which PM fails to run, and recovers known organogenesis dynamics, including gut convergence from visceral and definitive endoderm. CPM’s strengths are global hierarchy preservation, scalability, and biologically aligned geometry; we make these claims precise rather than asserting uniform superiority across every metric and dataset.

More broadly, CPM shows that contrastive representation learning and hyperbolic geometry are complementary: the former supplies scalable, preprocessing-free feature learning, and the latter supplies a space in which developmental hierarchies fit with low distortion. The resulting embeddings are not only compact but quantitatively interpretable, linking geometric position to developmental state and opening a path toward hierarchy-aware analysis of the increasingly large and multimodal datasets that now define single-cell biology.

## 2 Results

### 2.1 A single biologically aligned embedding streamlines single-cell analysis

Conventional scRNA-seq analysis proceeds through a long, multi-step pipeline. Raw counts pass through highly-variable-gene selection, PCA-based dimensionality reduction, batch-effect correction, clustering, a Euclidean visualization such as UMAP or t-SNE, and finally a separate trajectory- or pseudotime-inference step. Each stage introduces its own design choices and hyperparameters, and biases introduced early propagate downstream. CPM collapses this sequence into one model: it maps the count matrix directly to a two-dimensional hyperbolic embedding, requiring a single architectural design choice and one round of hyperparameter tuning, and removing the need for PCA, batch correction, clustering, and a dedicated trajectory-inference stage (Fig. 2). CPM also does not require highly-variable-gene (HVG) selection and can embed the full, unintegrated gene set directly; we applied HVG selection to some datasets only to follow their established pipelines and keep the comparison with baselines uniform.

**Figure 2.**
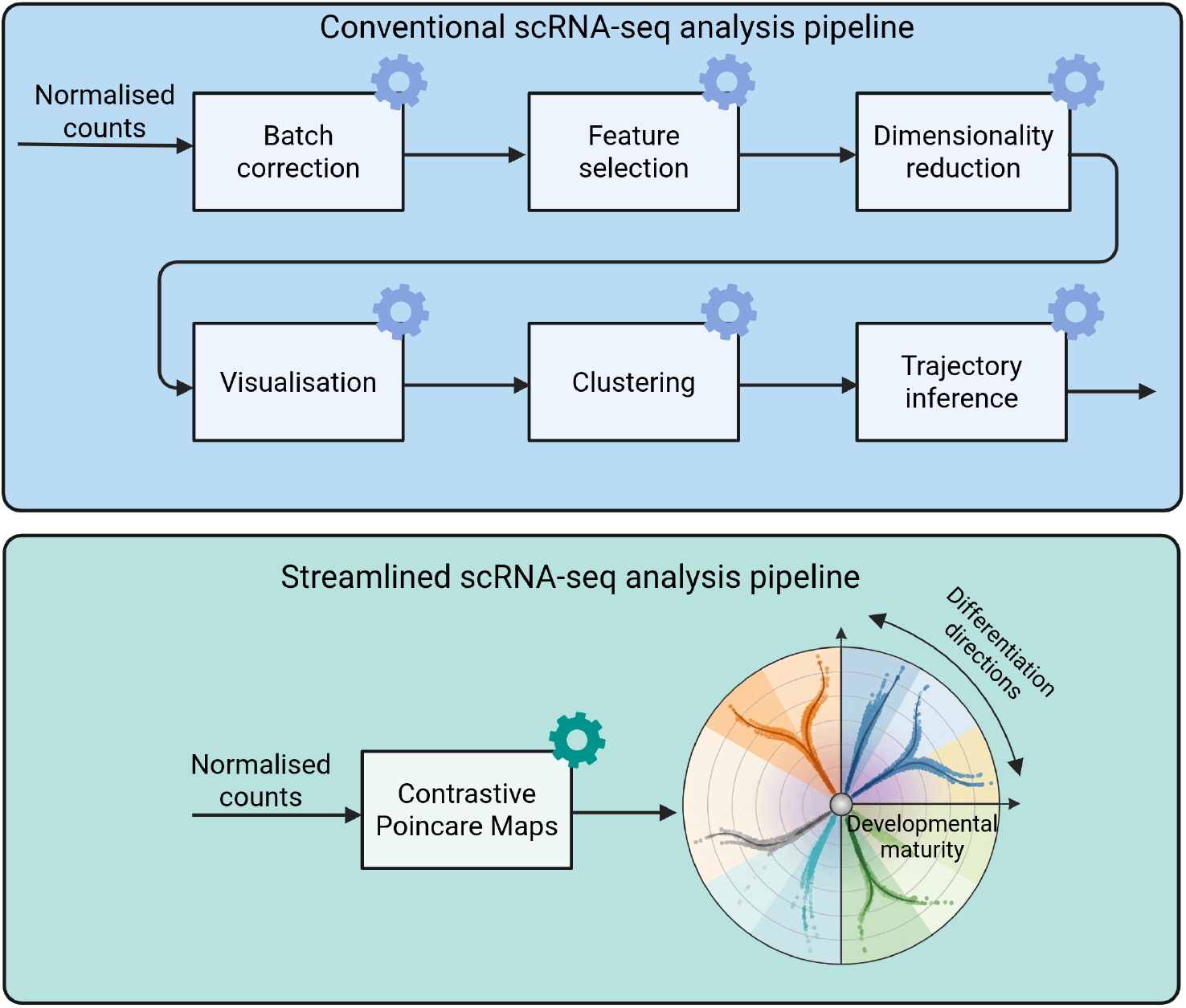
CPM replaces a multi-step analysis pipeline with a single, biologically aligned embedding. The conventional scRNA-seq pipeline: starting from normalized counts, highly-variable-gene selection, PCA-based dimensionality reduction, batch-effect correction, clustering, a Euclidean visualization (UMAP or t-SNE), and a separate trajectory- or pseudotime-inference step each introduce their own design choices and hyperparameters. The streamlined CPM pipeline: starting from the same normalized counts, a single architecture and one round of hyperparameter tuning map cells directly onto the two-dimensional Poincaré disk, in which radial distance from the root encodes developmental maturity and angular position encodes differentiation direction, so lineages fan out into distinct angular sectors and trajectory structure is read directly off the embedding geometry.

CPM takes the cells *×* genes count matrix and processes it in mini-batches. For each cell it builds a corrupted second view (12), encodes both views through a trainable projection block that replaces PCA and a hyperbolic block that places cells on the two-dimensional Poincaré disk (*κ* = *−*1) (9), and trains this encoder with an InfoNCE contrastive objective (10, 11) that draws similar cells together and pushes dissimilar cells apart in hyperbolic space (Methods). The two learned dimensions are biologically aligned rather than an arbitrary layout. Radial distance from the root encodes developmental maturity, and angular position encodes differentiation direction. This alignment lets a single embedding deliver analyses that conventionally demand separate tools: ordering cells by their radial distance recovers pseudotemporal progression comparable to diffusion pseudotime (16), while reading angular sectors recovers the canonical ordering of lineages. Trajectory structure is thus read directly off the embedding geometry, without a downstream inference step.

### 2.2 CPM preserves global hierarchy with low distortion at competitive cost

We benchmarked CPM against five baselines spanning two geometries: the hyperbolic Poincaré Maps (PM) (8), which fits precomputed pairwise similarities; the hyperbolic variational autoen-coder scPhere (13), trained by maximizing an evidence lower bound (ELBO) and the Euclidean methods t-SNE (5), UMAP (6), and PHATE (7). Benchmarking scPhere in its hyperboliclatent configuration isolates the effect of the learning objective from the effect of geometry: scPhere’s native Lorentz-model embedding converts isometrically to the same Poincaré disk used by CPM and PM, so only the learning objective differs, discriminative and contrastive for CPM, generative and reconstruction-based for scPhere. We quantified embedding quality with two complementary metrics, Q-scores (14) and worst-case distortion (Methods).

We built a synthetic benchmark with known ground truth: a balanced binary tree of depth five (63 branches), realized as 6,300 cells in a 1,000-dimensional feature space, where each branch is a 100-cell random walk seeded from its parent, so that each cell has a known branch of origin (Methods). Consistent with how CPM is intended to be used, all baselines received 50-component PCA-reduced input whereas CPM embedded the full 1,000-dimensional features directly. Reading a Poincaré-disk embedding differs from reading a Euclidean scatter plot: hierarchical depth is encoded radially, as distance from the origin, and separate branches occupy separate angular sectors. CPM produced the embedding that most closely matched the ground-truth tree, ordering successive generations along increasing radius and resolving branches into distinct angular sectors (Fig. 3a). The baselines distorted the hierarchy in characteristic ways: PM resolved only the first level and compressed all deeper generations onto the boundary of the disk (8); scPhere, within the same hyperbolic disk, recovered the tree less faithfully than CPM; UMAP introduced branch crossings that violate the hierarchy; t-SNE fragmented the tree into disconnected components; and PHATE preserved the coarse branching shape but with lower fidelity than CPM (Fig. 3a).

**Figure 3.**
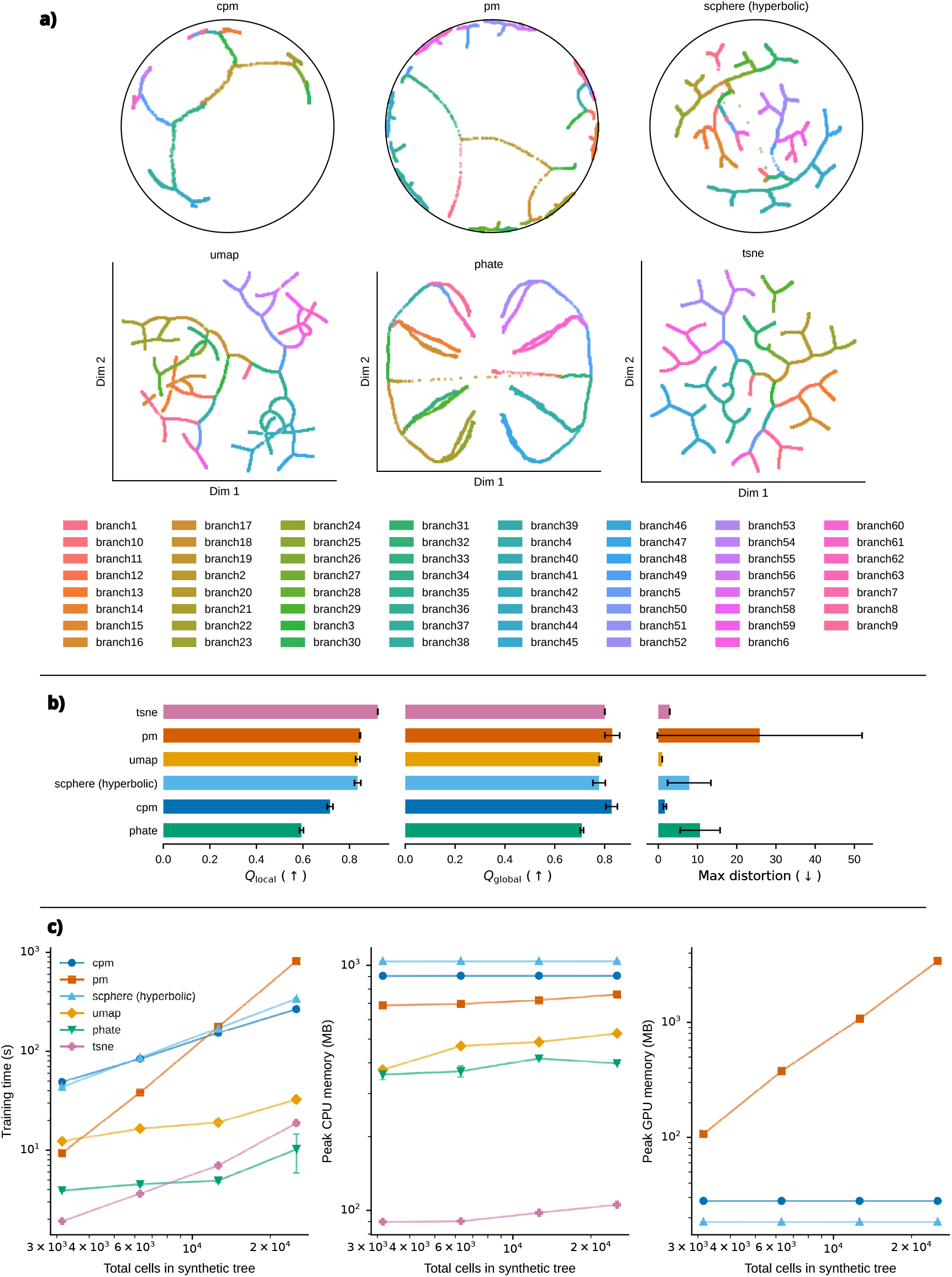
CPM combines faithful global hierarchy with low worst-case distortion and near-constant GPU memory on a synthetic benchmark. **a)** Embeddings of a depth-5 binary tree (63 branches, 6,300 cells) produced by CPM and the baselines. **b)** Local and global Q-scores and worst-case distortion for each method (mean *±* s.d. over 3 random seeds). **c)** Wall-clock training time and peak CPU and GPU memory as a function of dataset size (3,150-25,200 cells) for all baselines. PHATE, t-SNE and UMAP did not use the GPU.

These qualitative differences reflect a quantitative trade-off that no method escapes (Fig. 3b; mean *±* s.d. over three random seeds). Because local Q-scores, global Q-scores, and worst-case distortion reward different properties, no method scored best on all three. t-SNE attained the highest local Q-score (0.92 *±* 0.00) yet fragmented the tree, showing that strong local rank preservation need not yield a faithful global structure. PM achieved the best Q-scores among the hyperbolic methods (local 0.85 *±* 0.00, global 0.83 *±* 0.03) but incurred the highest worst-case distortion of any method (25.9 *±* 26.1). ScPhere achieved Q-scores comparable to UMAP (local 0.83 *±* 0.01, global 0.78 *±* 0.03) and held distortion far below PM’s (7.9 *±* 5.5 vs. 25.9 *±* 26.1), yet still trailed CPM on both distortion and global structure: within the same hyperbolic disk, the contrastive objective, not the geometry alone, accounts for CPM’s low distortion. UMAP achieved the lowest worst-case distortion (1.0 *±* 0.0) but preserved global structure least well among the embeddings (global Q 0.78 *±* 0.00) and crossed branches. CPM was the only method to pair top-tier global structure (global Q 0.83 *±* 0.02, on par with PM) with low worst-case distortion (1.7 *±* 0.4), far below PM (25.9 26.1), scPhere (7.9 *±* 5.5), and PHATE (10.7 *±* 5.1), and within twofold of the best Euclidean method (UMAP, 1.0 *±* 0.0). CPM’s distortion was also markedly more reproducible across seeds than either hyperbolic baseline. Its lower local Q-score (0.72 *±* 0.01) reflects a deliberate trade-off: the InfoNCE objective contrasts each cell against the full batch and so optimizes relative separation across all scales, rather than the exact ordering of nearest neighbors that the local Q-score rewards. CPM thus favors global hierarchy and worst-case fidelity, the properties that most directly govern reliable lineage detection, over fine local rank order.

CPM’s efficiency advantage lies in how it scales with dataset size. Across trees from 3,150 to 25,200 cells, CPM’s peak GPU memory stayed essentially constant at roughly 28 MB, because it trains on mini-batches, whereas PM materializes a pairwise similarity matrix with *O*(*n*^2^) memory complexity by design (8), from 0.11 GB to 3.4 GB over the same range, a more than 120-fold gap at the largest size (Fig. 3c). scPhere, also trained on mini-batches, likewise held GPU memory nearly constant (about 18 MB across all sizes). Peak CPU memory, by contrast, stayed within a comparable range for all three methods (roughly 690-1,050 MB) regardless of dataset size. Training time separates the two methods: CPM, trained on the full 1,000-dimensional features, and scPhere, trained on the 50-component PCA-reduced input, took comparable time at 3,150 cells (49 s vs. 44 s), but CPM scaled better as datasets grew, overtaking scPhere by 25,200 cells (267 s vs. 340 s). CPM’s training time also remained well below PM’s, overtaking it as datasets grew (267 s versus 817 s at 25,200 cells). CPM’s cost is thus dominated by a fixed training overhead rather than by dataset size, the regime that matters as single-cell datasets approach atlas scale.

In short, no single baseline matches CPM’s combination of properties: it preserves global hierarchy as faithfully as the best method, keeps worst-case distortion far below either hyperbolic alternative, and holds GPU memory nearly constant as datasets grow while training faster than scPhere at scale. This balance, not dominance on any one metric, is what suits CPM to large developmental datasets.

### 2.3 CPM’s geometry aligns with biological axes and recovers hematopoietic lineage branching

Unlike the synthetic benchmark, real developmental data carry no ground-truth tree. We therefore tested CPM against mouse hematopoiesis, a lineage hierarchy established independently of any embedding method. Paul et al. profiled 2,730 mouse bone marrow myeloid progenitor cells by MARS-seq and established that common myeloid progenitors split into two committed branches, the megakaryocyte-erythrocyte progenitor (MEP) lineage and the granulocyte-monocyte progenitor (GMP) lineage (15). We used the same 1,000 highly variable genes selected by the “zheng17” preprocessing recipe (18) as Klimovskaia et al. (8).

The CPM embedding of this dataset makes the geometry-to-biology correspondence explicit: distance from the root marks developmental maturity and angular position marks differentiation direction (Fig. 4a). To test whether these coordinates reconstruct the known branching order without an additional clustering or trajectory-inference step, we plotted each cell type’s median distance from the root against its angular position. CPM reproduces the established order: the MEP lineage, comprising the uncommitted MEP progenitor pool together with its erythroid and megakaryocyte descendants, and the GMP lineage resolve into distinct angular sectors that diverge from the root at comparable radius (Fig. 4a).

**Figure 4.**
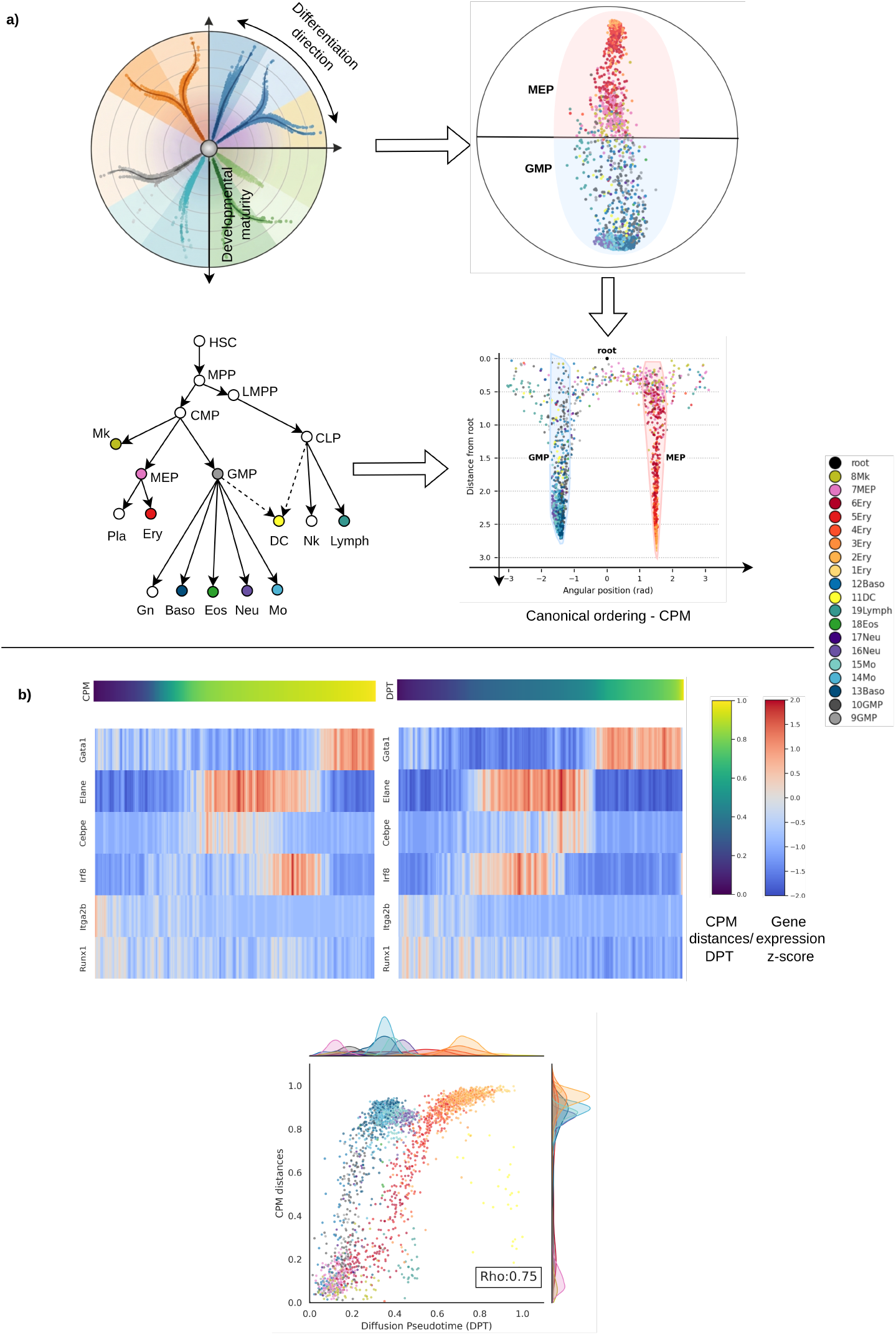
CPM’s geometry recovers hematopoietic lineage branching and pseudotemporal ordering. **a)** A guide to reading the Poincaré disk (as in Fig. 2), the CPM embedding of mouse hematopoiesis marked by MEP and GMP lineage, median distance from root plotted against angular position showing the MEP and GMP lineages bifurcating into distinct angular sectors, and the established canonical hematopoietic lineage tree for reference. **b)** Cells ordered by CPM root distance and by diffusion pseudotime (DPT), with expression trends for Gata1, Elane, Cebpe, Irf8, Itga2b, and Runx1, and a scatter plot of CPM distance against DPT value, colored by cell type, with the Spearman correlation coefficient (*ρ* = 0.75) annotated.

Ordering cells by CPM root distance reproduces the expression trends that diffusion pseudotime (DPT) (16) assigns to key lineage markers, Gata1 and Itga2b (MEP, erythroid and megakaryocyte respectively), Elane and Cebpe (GMP, granulocyte), Irf8 (the dendritic sub-lineage within GMP), and Runx1, and CPM distances correlate with DPT values (Spearman *ρ* = 0.75; Fig. 4b). A single CPM embedding thus recovers both the branching topology and the pseudotemporal ordering that conventional analysis obtains from two separate tools.

### 2.4 CPM scales to a 116,000-cell organogenesis atlas and recovers its differentiation dynamics

Mouse hematopoiesis has a single progenitor and two committed branches, a comparatively simple topology. We tested CPM on a larger, multi-lineage dataset with a more complex, non-linear topology to determine whether its global-hierarchy preservation and memory advantage generalize. We embedded the mouse gastrulation and early organogenesis atlas (17), 116,312 cells collected across nine stages, from embryonic day 6.5 to embryonic day 8.5 (E6.5-E8.5), spanning epiblast, primitive streak, and mesodermal, endodermal, ectodermal, and haematoendothelial derivatives. Following the original study, we normalized counts to 10,000 per cell, applied a log1p transform, and retained 2,000 highly variable genes; we did not apply PCA, batch correction, or clustering before training (Methods).

At this scale, PM’s quadratic memory requirement becomes prohibitive rather than merely slow: computing the pairwise similarity matrix for all 116,312 cells exhausts memory on a cluster with 112 CPUs and 1 TB of RAM, so PM does not produce an embedding for the full dataset. CPM trains on mini-batches and completes in approximately 2 to 3 hours on the same data, carrying the memory advantage established on synthetic trees (Fig. 3c) over to a real, atlas-scale dataset.

Similarly, the hyperbolic variational autoencoder scPhere encounters strict numerical limits as heterogeneity and dataset size increase further. Even on a 58,156-cell subsample of this dataset, the high variance forces the wrapped-normal *µ* to grow unboundedly over training, eventually overflowing the hyperbolic trigonometric functions. While tightening the internal boundary clamp or adding a quadratic radius penalty to the ELBO loss can offer temporary fixes for a few narrow hyperparameter choices, these often render the Monte-Carlo KL estimate degenerate at the boundary, causing the optimizer to exploit the artifact and the loss to diverge to *−*10^12^. Training the contrastive objective of CPM, on the other hand, remained numerically stable for the entire 116,312-cell dataset throughout: embeddings did not collapse or diverge near the boundary of the disk.

The preceding sections examined a single cell’s radial distance and angular position relative to the embedding’s root. Here we turn to lineage-scale relationships between arbitrary pairs of cells, read from the geodesic connecting them, the shortest path between two points in the Poincaré disk. Fig. 5a,b shows the full CPM embedding, colored by lineage and by embryonic stage. From a reference epiblast/primitive-streak cell (star), the geodesic angle to a given cell tracks differentiation direction and matches the lineage coloring in (a), while geodesic distance to that cell tracks developmental maturity and matches the stage coloring in (b): for example, the geodesic connecting an E7.0 epiblast cell to an E8.5 cell shifts in angle as well as length, tracing the intervening change in fate together with the intervening developmental time. Fig. 5c decomposes this global view stage by stage. At each stage we mark that stage’s biological progenitor along the epiblast to primitive streak to neuromesodermal progenitor (NMP) sequence and draw its geodesics to the fates the stage gives rise to. Cells move away from the progenitor with developmental time and separate by lineage, consistent with (a,b); at E6.75 to E7.75 the progenitor sits close to the E6.5 epiblast cell because epiblast, primitive streak, early mesoderm, and early endoderm genuinely co-locate in a low-radius hub at this stage, reflecting their shared immature state.

**Figure 5.**
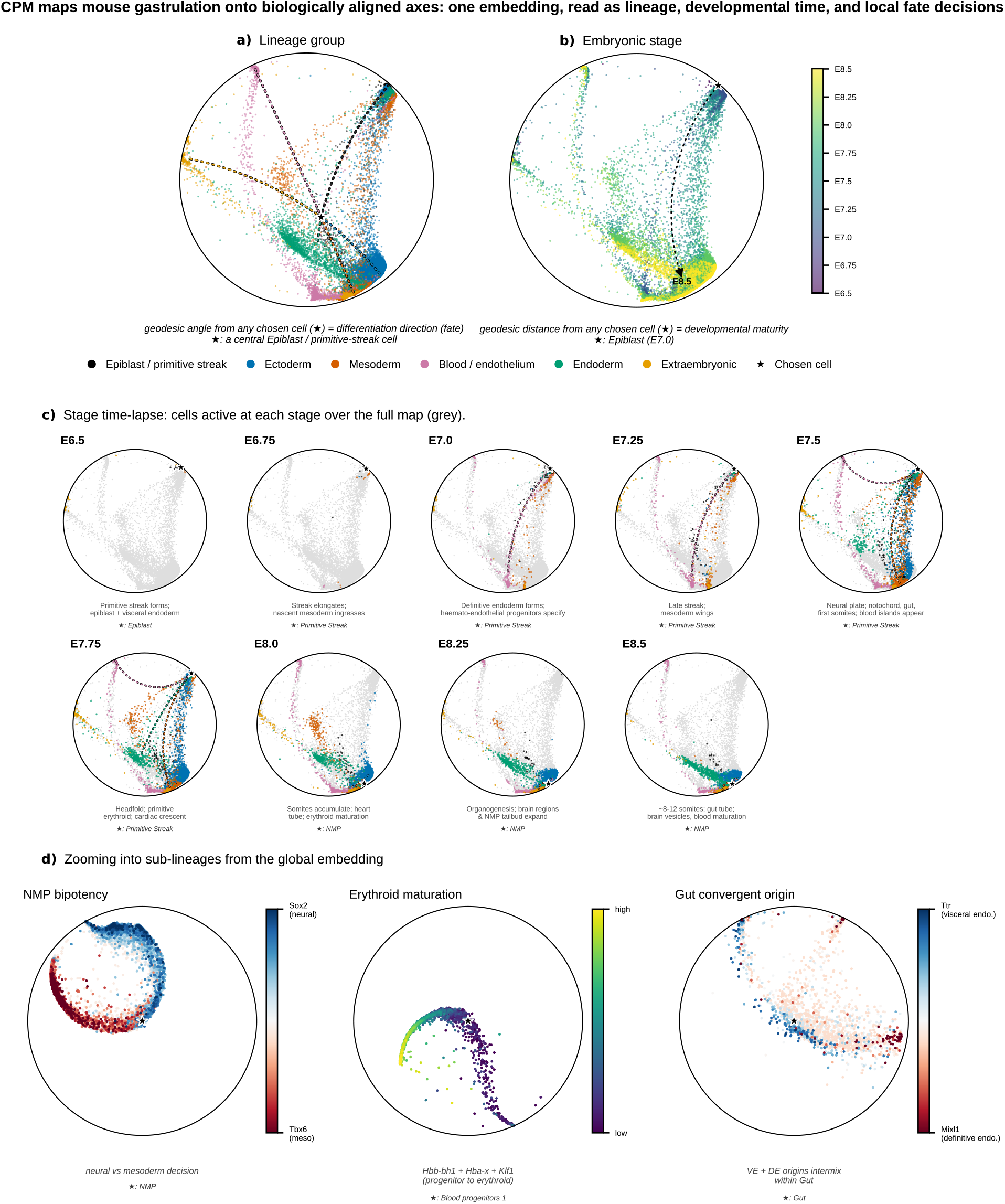
CPM’s geometry recovers organogenesis dynamics across the 116,312-cell mouse gastrulation atlas. Star, reference cell; lines, geodesics. **a**,**b)** Embedding colored by lineage (a) and stage (b); geodesic angle from the star tracks differentiation direction, distance tracks maturity. **c)** Stage time-lapse (E6.5-E8.5); star marks each stage’s progenitor, with geodesics to its fates. **d)** Sub-lineages re-rooted at their progenitors: NMP bipotency (Sox2 minus Tbx6), erythroid maturation (Hbb-bh1/Hba-x/Klf1), gut convergence (Ttr minus Mixl1).

A property specific to hyperbolic embeddings is that an isometric Möbius translation re-centers any cell at the disk origin without retraining and without distorting local distances, so a sub-lineage can be zoomed into while keeping the geometry that governs the rest of the analysis, a capability Euclidean embeddings lack. Fig. 5d applies this re-rooting to three sub-lineages that illustrate complementary local topologies, showing in each case only the cells belonging to that sub-lineage. Re-rooted at the NMP progenitor, a Sox2 (neural) minus Tbx6 (mesoderm) score separates into two clean arcs, recovering NMP bipotency as a divergence. Rerooted at blood progenitors, a composite erythroid score (mean z-scored Hbb-bh1, Hba-x, Klf1) increases monotonically along a single arc, recovering erythroid maturation as a progression. Re-rooted at the gut cell type, a visceral-endoderm score (Ttr) minus a definitive-endoderm score (Mixl1) places visceral-endoderm-derived and definitive-endoderm-derived cells at opposite poles that converge on gut, recovering the convergent origin of gut endoderm from both visceral and definitive endoderm reported in (17), directly from the embedding and without a separate trajectory-inference step.

Together, these results show that CPM’s global-hierarchy preservation and memory advantage, established on synthetic trees, carry over to a real atlas-scale dataset on which the hyperbolic baseline cannot run at all, and that its geometry recovers known organogenesis biology, a lineage bifurcation (NMP), a linear maturation trajectory (erythropoiesis), and a convergent origin (gut endoderm), directly and without additional processing.

## 3 Discussion

Euclidean embeddings cannot accommodate the exponential growth of a developmental tree without distorting it; hyperbolic space can, but PM, one of the existing hyperbolic methods for single-cell data, reaches that geometry through a transductive, hand-engineered pipeline, precomputed pairwise similarities, estimated manifold metrics, and PCA preprocessing, that scales quadratically in memory and must be rebuilt for every dataset. Likewise, previous attempts to learn hyperbolic embeddings via variational autoencoders (e.g., scPhere) suffer from severe numerical instability at scale, as the wrapped-normal distribution overflows hyperbolic trigonometric functions and boundary clamps collapse the KL divergence estimate.

CPM replaces this pipeline with a learned contrastive encoder that maps raw expression directly onto the Poincaré disk, dispensing with PCA, batch correction, and clustering not only for efficiency but because each of these steps makes its own choices and can discard or distort signal before the embedding ever sees the data. This substitution is what makes the rest of the paper’s results possible: near-constant memory as datasets grow, low worst-case distortion alongside faithful global hierarchy on a synthetic tree with known ground truth, and, on two independent developmental datasets, a two-dimensional embedding whose radial and angular coordinates recover established biology, MEP/GMP branching and DPT-correlated pseudotime in mouse hematopoiesis, and NMP bipotency, erythroid maturation, and VE/DE gut convergence in a 116,312-cell gastrulation atlas, directly and without a separate trajectory-inference step, at a scale where PM does not run at all.

This advantage is not uniform across every property an embedding can be judged on, however. CPM’s contrastive objective contrasts each cell against its full batch, which optimizes global hierarchy and worst-case distortion at some cost to local rank order: on the synthetic benchmark, methods such as t-SNE that are not built for hierarchy preservation nonetheless attain a higher local Q-score. For shallow trees or small, single-progenitor datasets, this trade-off matters less, and simpler methods can be competitive. CPM’s design instead targets the regime where it matters most: deep, multi-lineage, atlas-scale developmental data, where a hand-engineered, quadratic-memory pipeline is no longer an option and global hierarchy is the property lineage detection depends on.

This same global-first geometry is also what makes the embedding quantitatively interpretable: radial position, angular separation, and geodesics between arbitrary cells all carry consistent geometric meaning throughout the disk, so comparisons across lineages and sub-lineages are possible in a way an arbitrary Euclidean layout does not support. Reading these relationships in practice, as in the gastrulation analysis, requires choosing a reference cell for each sub-lineage; this is a deliberate, interpretable design choice rather than an automated step, and developing principled, data-driven reference selection is a natural extension.

More broadly, nothing in CPM’s architecture is specific to gene expression. The model consumes a matrix of samples by features, forms positive pairs by corrupting feature entries, and optimizes a contrastive objective against hyperbolic distance; the biology enters only through what the features mean. The same recipe therefore applies wherever data carry a latent hierarchy whose states outgrow what a Euclidean layout can hold: taxonomies, ontologies, citation and knowledge graphs, and hierarchical structure in text. Within single-cell biology, the natural extensions are spatial transcriptomics, chromatin accessibility, and multimodal integration across these modalities. We validated CPM on developmental transcriptomics alone, so its behavior on other data types remains to be established. What generalizes with confidence is the underlying observation: a discriminative objective and a negatively curved space address complementary halves of one problem, scalable representation learning and low-distortion accommodation of exponential growth. As datasets grow in any of these domains, embeddings that scale linearly in memory and remain interpretable by construction offer a path toward hierarchy-aware analysis that does not trade off fidelity for size.

## 4 Methods

Let 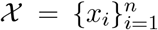 be the given scRNA-seq dataset with *n* cells and *d* features per cell, i.e., *x*_*i*_ *∈* ℝ^*d*^. In this work, we focus on representing data in hyperbolic space by learning a mapping, *ϕ* : *X →* ℙ^2^ between data space *X* and hyperbolic representation space P^2^, which maps a data point *x*_*i*_ *∈ X* to *ϕ*(*x*_*i*_) *∈* ℙ^2^.

### 4.1 Hyperbolic geometry

Hyperbolic spaces are smooth Riemannian manifolds with constant negative curvature. We adopt the Poincaré ball model, which defines a hyperbolic space within the Euclidean unit ball, i.e.,

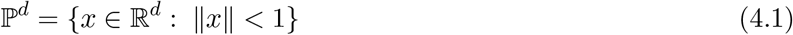

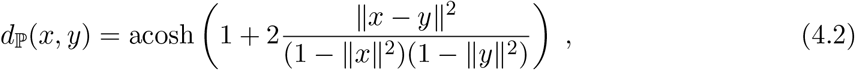

where *l/ · l/* denotes the usual Euclidean norm. The exponential map Exp_*x*_ : *T*_*x*_ℙ^*d*^ *→* ℙ^*d*^, an important geometric tool that characterises geodesics in the space, is given in terms of Möbius addition (3, 9) by

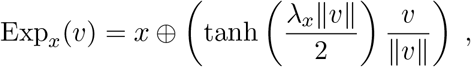

where 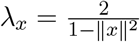 is the conformal factor at *x* and

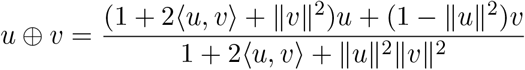

is Möbius addition. At the origin this reduces to Exp 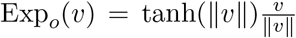, the map used to place the encoder’s output onto the disk (Model architecture and optimization).

Geodesics in the Poincaré model can be computed via simple retractions of the form

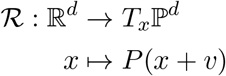

where *P* is an orthogonal projection onto ℙ^*d*^, given by

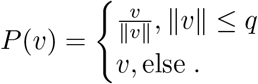

Throughout this work, we focus on the two-dimensional setting, ℙ^2^.

### 4.2 Model architecture and optimization

CPM adopts the sampling strategy of SCARF (12). For each batch, a positive sample is obtained by constructing an additional corrupted view: a randomly chosen fraction of feature entries is substituted with values from other training examples. The encoder consists of a projection block and a hyperbolic block. The projection block is composed of linear layers with ReLU activation and performs both dimensionality reduction and noise reduction, adapted and trained end-to-end unlike PCA. The hyperbolic block comprises hyperbolic linear layers with ReLU activation (9). We use a two-dimensional representation space, the Poincaré disk, with curvature *κ* = *−*1. A single forward pass of the model can be written as

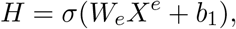

where *H* is the latent representation learnt by the projection block, *W*_*e*_ are the parameters of the projection block, *X*^*e*^ is the input data, *b*_1_ is the bias, and *σ*(.) is the ReLU activation. The representation is then projected onto the Poincaré disk using the exponential map Exp_*o*_(.) and processed by the hyperbolic block:

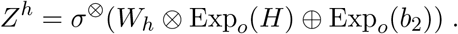

Here, *W*_*h*_ and *b*_2_ are the trainable parameters of the hyperbolic block and *Z*^*h*^ is the hyperbolic representation learnt by the model; *σ*^*⊗*^(.) = Exp_*o*_(*σ*(Log_*o*_(.))), where Log_*o*_(.) is the logarithmic map (9).

Hyperbolic representations of both augmented views are learnt using encoder blocks with shared weights. Their similarity, the *Poincaré similarity*, is computed from the Poincaré distance (Eq. 4.1) as 1*/*(1 + *d*_P_(*x, y*)). These similarities enter the classical InfoNCE contrastive loss for training. We use Riemannian Adam (19) for optimization, with different learning rates for the projection and hyperbolic blocks.

### 4.3 Synthetic data generation

We generated synthetic datasets using PHATE (7), which relies on diffusion-limited aggregation. Ground-truth hierarchies were balanced binary trees, with each depth level branching into two subtrees, realized as a cumulative random walk of independent uniform increments per branch, each child branch initiated from its parent’s terminal point, with isotropic Gaussian noise added to the completed data matrix.

### 4.4 Data pre-processing

The synthetic data was standardized to zero mean and unit variance. The mouse hematopoiesis dataset was obtained pre-processed from Klimovskaia et al. (8), using the “zheng17” recipe from Scanpy (18), which selects 1,000 highly variable genes across 2,730 cells; we follow this choice for direct comparability with PM, whose own preprocessing it is. The mouse gastrulation dataset was normalized to 10,000 counts per cell, log1p-transformed, and filtered to 2,000 highly variable genes, following the gene selection used by Pijuan-Sala et al. (17) in the original study. While CPM operates without additional dimensionalitythe metric’s known dependence reduction, PCA was applied to all baselines along with their respective standard pre-processing steps.

### 4.5 Hyperparameter tuning and reproducibility

The most important hyperparameters are batch size, corruption rate for data augmentation, the number of projection and hyperbolic layers, latent dimension, and the learning rates for the projection and hyperbolic blocks; the hyperbolic block’s learning rate is usually higher than the projection block’s. A batch size of 128 worked well for most datasets in this study (batch size sweep in Supplementary Fig. 6). The number of hyperbolic layers depends on the depth of the input hierarchy; we used 2 hyperbolic layers for the deepest synthetic tree. Balancing the number of projection layers against the number of cells considered for augmentation in a batch controls embedding distortion; a corruption rate of 0.2 with one projection layer achieved good performance on most datasets.

### 4.6 Complexity

The memory complexity of CPM is *O* (*b*^2^), where *b* is the batch size, dominated by the pairwise similarity calculation within each batch used for the contrastive loss; the time complexity of this calculation is likewise *O* (*b*^2^), and for the full training process *O* (*b*^2^*ed*), where *e* is the number of epochs and *d* is the feature dimension. Training loss converged within 200 epochs for all datasets in this study. Training time ranged from under 15 minutes for synthetic datasets to 2-3 hours for the mouse gastrulation dataset (*>*100,000 cells and 2,000 genes); total memory consumption stayed below 20GB throughout.

### 4.7 Quantitative evaluation metrics

Let *ϕ* : *X → ℳ* denote an embedding of a dataset *X*. For CPM and PM, *ℳ* = ℙ^2^, the hyperbolic plane; for Euclidean embeddings, *ℳ* = ℝ^*d*^ for small *d*.

#### Distortion

Consistent with conventions in the classical metric embedding literature, the representation error of *ϕ* can be quantified using *distortion*:

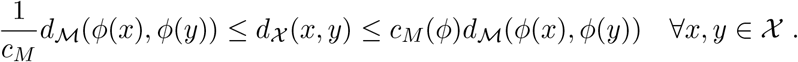

If *ϕ* is an isometric embedding, *c*_*M*_ (*ϕ*) = 1.

#### Worst-case distortion

The parameter *c*_*M*_ can be difficult to compute in practice, as it requires access to *ϕ* in closed form over the whole data space. We therefore report the worst-case computational distortion score (see, e.g., (20)), which is most relevant when robustness guarantees are required across all pairs of points:

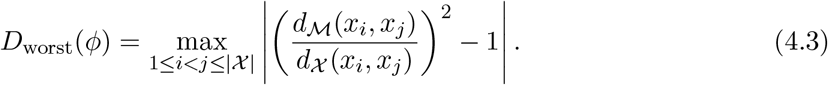

Here, *d*_*ℳ*_ denotes distance in the representation space (hyperbolic or Euclidean) and *d*_*X*_ distance in the data space. If *ϕ* is an isometric embedding, *D*_worst_ = 0.

#### Q-scores

Q-scores, introduced by Lee et al. (14), are widely used in the manifold embedding literature. Let *Q* = *{q*_*kl*_*}*_*k,l∈*[*N*]_ be the *co-ranking matrix* defined as

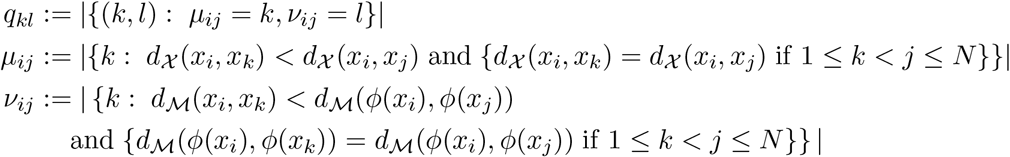

Here, |*I*| denotes the cardinality of a set *I*. The Q-curves are then given by

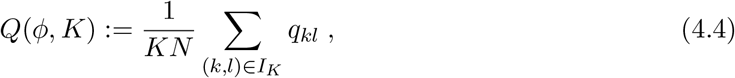

computed over blocks *I*_*K*_ := *{*1, …, *K} × {*1, …, *K}* of the co-ranking matrix, restricted to the upper half of *I*_*K*_ by symmetry. The local and global Q-scores of embedding *ϕ* are computed as

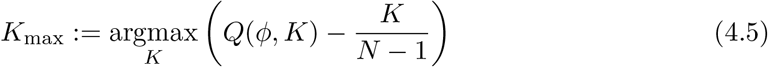

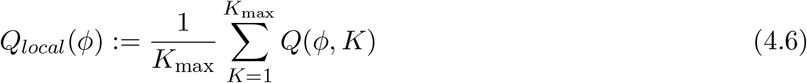

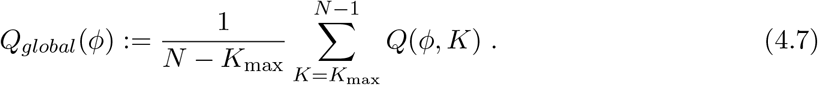

The split of the Q-curve at *K*_max_ distinguishes the local from the global regime. *Q*_*local*_ and *Q*_*global*_ range from 0 (worst) to 1 (best).

### 4.8 Geometric reading of the embedding

Contrastive Poincaré Maps are visualized using the 2D Poincaré disk layout (8). By default, the point with the smallest average distance to all others is placed at the origin. Alternatively, the embedding can be translated so that a chosen reference point *r* is mapped to the origin, while preserving all pairwise hyperbolic distances:

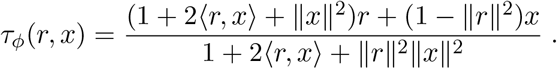

Owing to the increased resolution near the origin in hyperbolic space, this re-centering is used for inspecting hierarchical structure around a known root and for isolating sub-lineages in multi-lineage datasets (Fig. 5d).

A direct geodesic arc between two arbitrary points *p, q ∈* ℙ^2^ is a circular arc orthogonal to the disk boundary and is awkward to parametrize directly. We instead compute each geodesic independently: (i) apply *τ*_*ϕ*_(*−p, ·*), the Möbius isometry that sends *p* to the origin, giving the image *q*_*p*_ of *q* in this frame; (ii) sample a straight radial line from the origin to *q*_*p*_, with points spaced by tanh to give correct hyperbolic arc length; (iii) apply the inverse translation *τ*_*ϕ*_(*p, ·*) to map the sampled line back into the original coordinates. Because Möbius transformations are isometries of the hyperbolic metric, they map geodesics to geodesics, so the straight line computed in the *p*-centered frame maps back to the true geodesic between *p* and *q*. Each geodesic shown (root to group centroid, progenitor to fate) is computed by its own call to this procedure and does not persistently transform the embedding.

## 4.9 Marker-gene composite scores

For the sub-lineage analyses in Fig. 5d, each gene was z-scored within the cell subset shown in that panel, and composite scores were formed by averaging or differencing the relevant z-scored genes: the NMP score is Sox2 minus Tbx6; the erythroid score is the mean of z-scored Hbb-bh1, Hba-x, and Klf1; the gut-convergence score is Ttr minus Mixl1.

### 4.10 Baselines

CPM was benchmarked against PM (8), the hyperbolic variational autoencoder scPhere in its hyperbolic-latent configuration, and the Euclidean methods t-SNE (5) (Scikit-learn implementation (21), Barnes-Hut approximation), UMAP (6), and PHATE (7). All baselines received PCA-reduced input (Data pre-processing); CPM did not.

## Acknowledgements

The authors would like to thank Alice Bizeul for her input during the initial phase of the project.

## Author Contributions

Conceptualization: M.W., N.B.; Methodology: M.W., N.B.; Investigation & Formal Analysis: N.B., L.B., V.K.; Validation: M.W., H.C., N.B., L.B.; Visualization: N.B.; Writing and review: N.B., M.W., L.B., V.K. wrote the initial manuscript. All authors reviewed and edited the manuscript; Resources and funding: S.S., M.W.; Supervision: M.W.

## Use of Large Language Models and other software

The authors used a large language model (Claude, Anthropic) to assist with drafting and editing prose based on the authors’ own experimental design, data, and analyses. The model was not used to independently generate scientific claims or interpret results. Figure 2 was created using BioRender. The example Poincare disk in Figures 2 and 4 (showing developmental maturity and differentiation directions) was generated by Gemini AI.

## Data and Code Availability

A diverse collection of synthetic datasets were generated, following diffusion limited aggregation method described in Moon et al. (7) (https://github.com/KrishnaswamyLab/PHATE). The public datasets used in this study were: 1. Mouse gastrulation and early organogenesis (17) (https://github.com/MarioniLab/EmbryoTimecourse2018), 2. Mouse hematopoeisis (15) (processed data from: (8) https://github.com/facebookresearch/PoincareMaps).

The code is available at https://github.com/NithyaBhasker/ContrastivePoincareMaps.git.

## Additional results: Model robustness

CPM maintains stable performance across a broad range of hyperparameter settings (Supplementary Fig. 6). The number of projection layers and the corruption rate require a moderate balance to minimize distortion; outside this range, performance degrades gradually rather than catastrophically.

We further assessed whether CPM embeddings reflect stable manifold structure or vary as an artifact of training stochasticity, holding the input data and all hyperparameters fixed and varying only the random seed. We trained CPM from five fixed seeds (42, 123, 456, 789, 1234) on a synthetic depth-3 balanced binary tree (15 branches, 100 cells per branch; 1,500 cells, 1,000 features), scoring each resulting embedding independently against the raw 1,000-dimensional reference space (Fig. 7).

**Figure 6.**
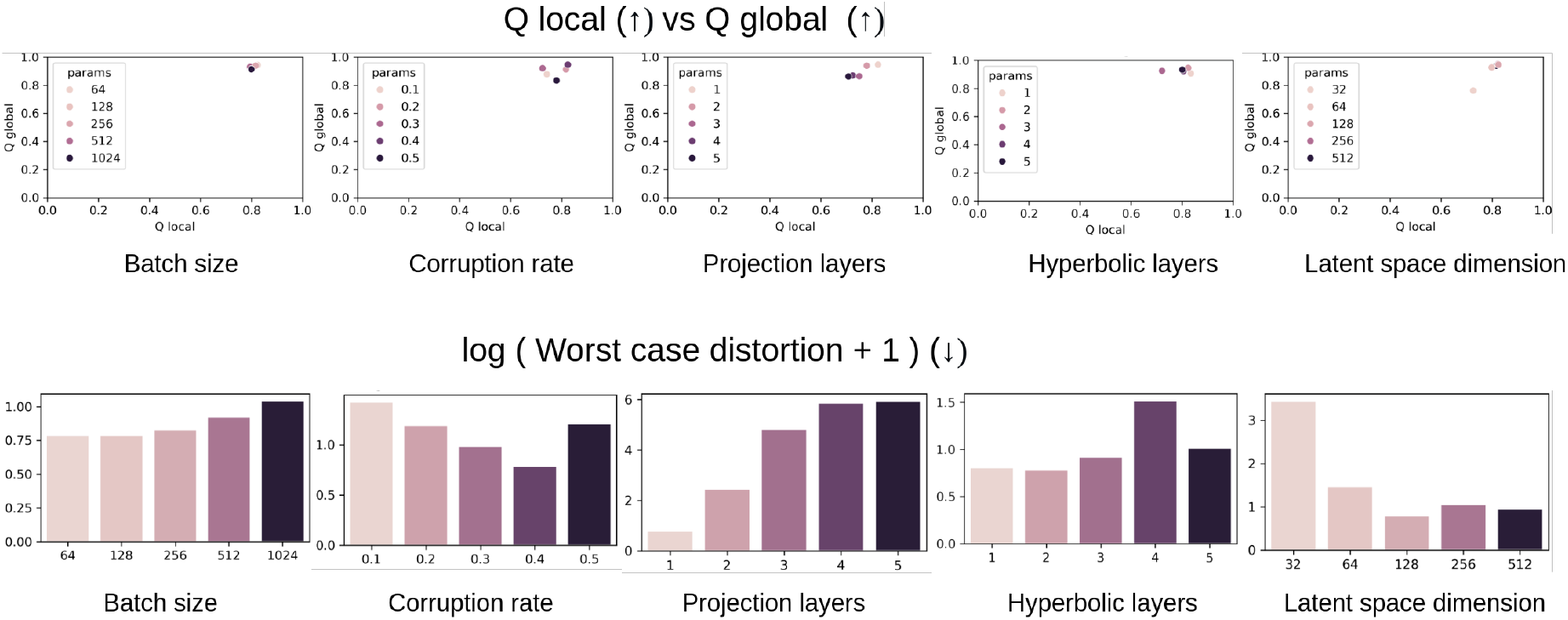
Robustness of CPM to hyperparameter variation. Quality metrics for CPM across a range of hyperparameter configurations (projection layers, corruption rate, batch size, latent dimension).

**Figure 7.**
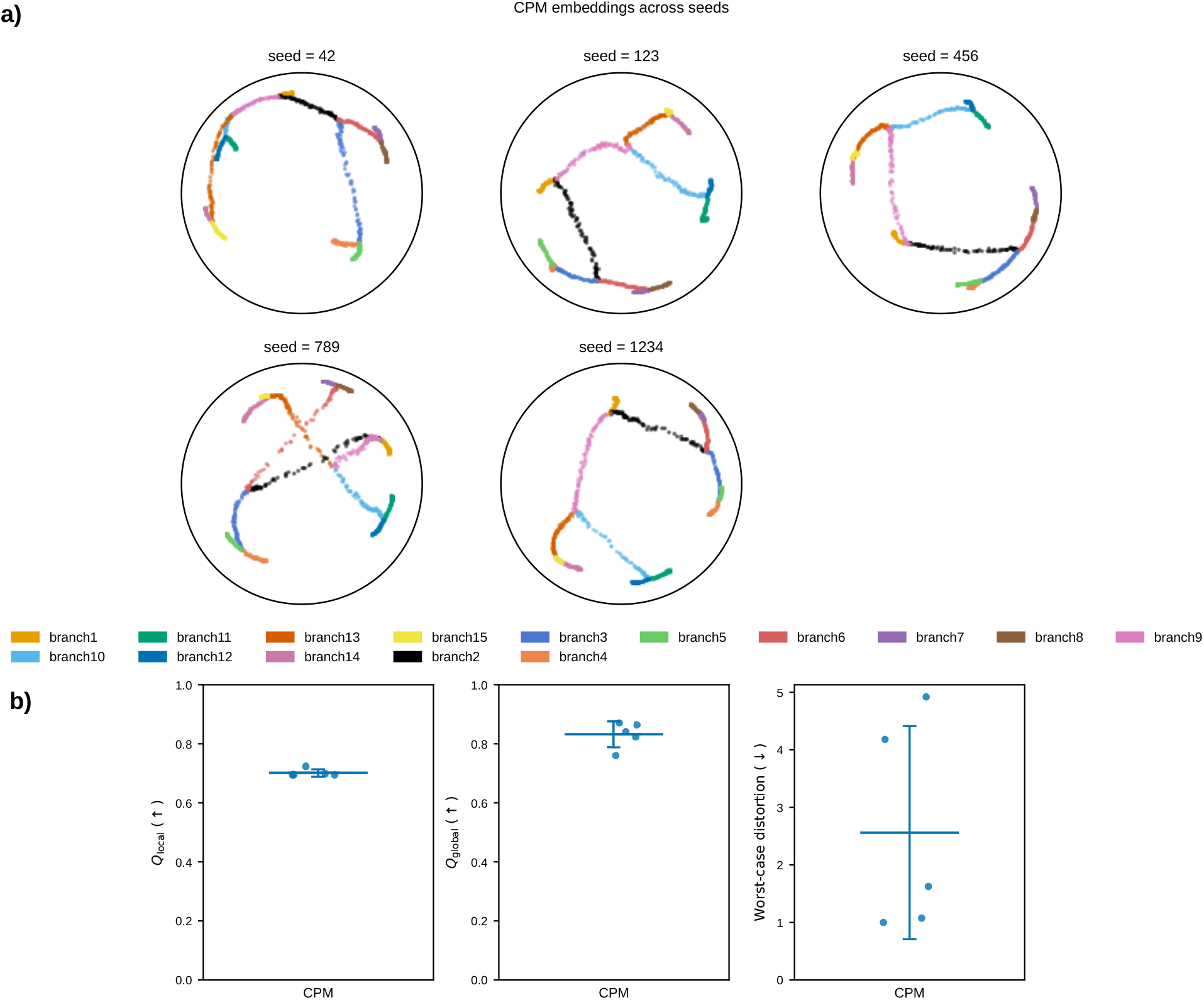
Run-to-run reproducibility of CPM across five random seeds. **a)** CPM embeddings of a synthetic depth-3 tree (1,500 cells), trained from five fixed seeds with all other hyperparameters and the input data held fixed, colored by ground-truth branch identity. **b)** Quality metrics for each seed.

Local neighborhood preservation (*Q*_*local*_) is highly reproducible across seeds (coefficient of variation 1.7%). Global structure preservation (*Q*_*global*_) is reasonably stable (CV 5.3%), with one seed (789) a mild low outlier. Worst-case distortion is markedly more seed-sensitive (CV 72%), separating into two seeds with elevated distortion (4.18-4.92) and three seeds near the metric’s practical floor (1.00-1.62). This reflects the metric’s known dependence on a single extreme pairwise distance rather than a change in aggregate embedding quality: the larger run-to-run variance in worst-case distortion is expected behavior for this metric, not evidence that the embedding itself is unstable.

We additionally assessed sensitivity to two architectural choices fixed elsewhere in this study, curvature and embedding dimension, training CPM from a single seed on the same synthetic depth-3 tree (1,500 cells) used for the reproducibility experiment above, with all other hyperparameters held fixed.

Across curvature magnitudes *κ∈ {−*0.5, *−*1.0, *−*2.0*}* (Supplementary Fig. 8), embedding quality is essentially flat: *Q*_*local*_ and *Q*_*global*_ vary by at most 0.4%, and worst-case distortion by about 2%, all within the seed-to-seed variability already characterized above (CV 1.7-72%). This spread falls within the seed-to-seed noise floor established above (CV 1.7-72%), so any genuine curvature effect in this range is small relative to ordinary training stochasticity and does not change the practical conclusion: curvature *κ* = *−*1, used throughout this study, is a reasonable default rather than a finely tuned choice.

Embedding dimension, by contrast, has a clear, consistent effect that exceeds seed noise (Supplementary Fig. 9). Increasing from *d* = 2 to *d* = 3 produces a sizeable gain in local structure preservation (*Q*_*local*_: 0.700 to 0.782) and roughly halves worst-case distortion (4.92 to 2.46); further increases to *d* = 4 and *d* = 5 continue to reduce worst-case distortion monotonically, to 1.36 at *d* = 5, while *Q*_*local*_ and *Q*_*global*_ plateau or fluctuate mildly around their *d* = 3 values. This matches the expected geometric picture: a two-dimensional hyperbolic disk has limited capacity to embed all branches of a 15-branch tree without forcing some pairs into worse relative distances, and additional dimensions relieve this packing constraint, with the largest gain from *d* = 2 to *d* = 3. We nonetheless use *d* = 2 throughout this study, both for direct comparability with the Euclidean and hyperbolic baselines, which are likewise evaluated in two dimensions, and because two-dimensional embeddings remain the standard for visualization and interpretability in single-cell analysis; CPM’s architecture extends directly to higher *d* when visualization is not the goal.

**Figure 8.**
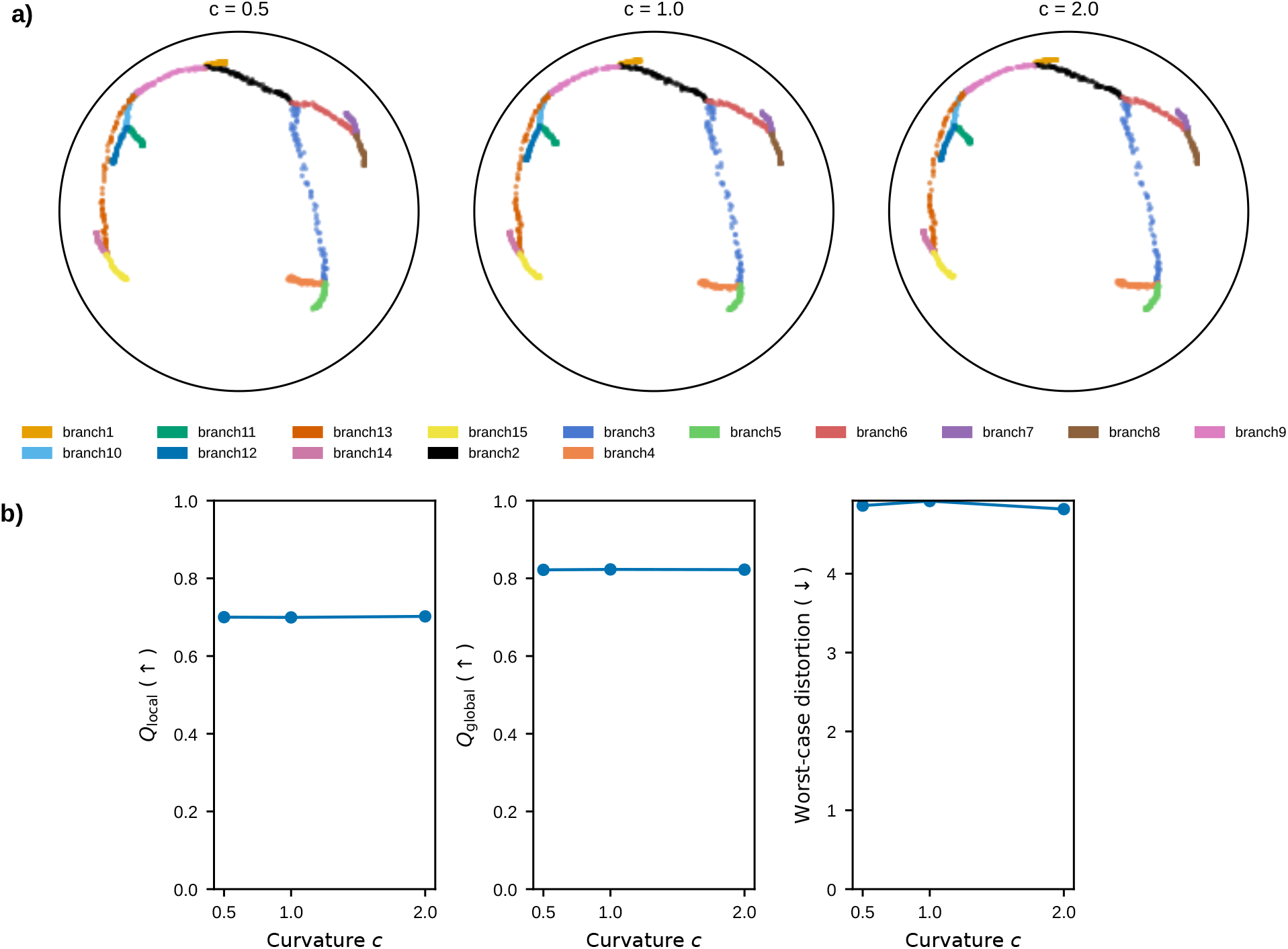
Sensitivity of CPM to curvature. **a)** CPM embeddings of the synthetic depth-3 tree in the two-dimensional Poincaré disc across curvature *κ ∈ {−*0.5, *−* 1.0, *−* 2.0*}*. **b)** Quality metrics (*Q*_*local*_, *Q*_*global*_, worst-case distortion) across the full curvature sweep, single seed, all other hyperparameters held fixed. Metrics vary only within the range of seed-to-seed noise characterized in Fig. 7.

**Figure 9.**
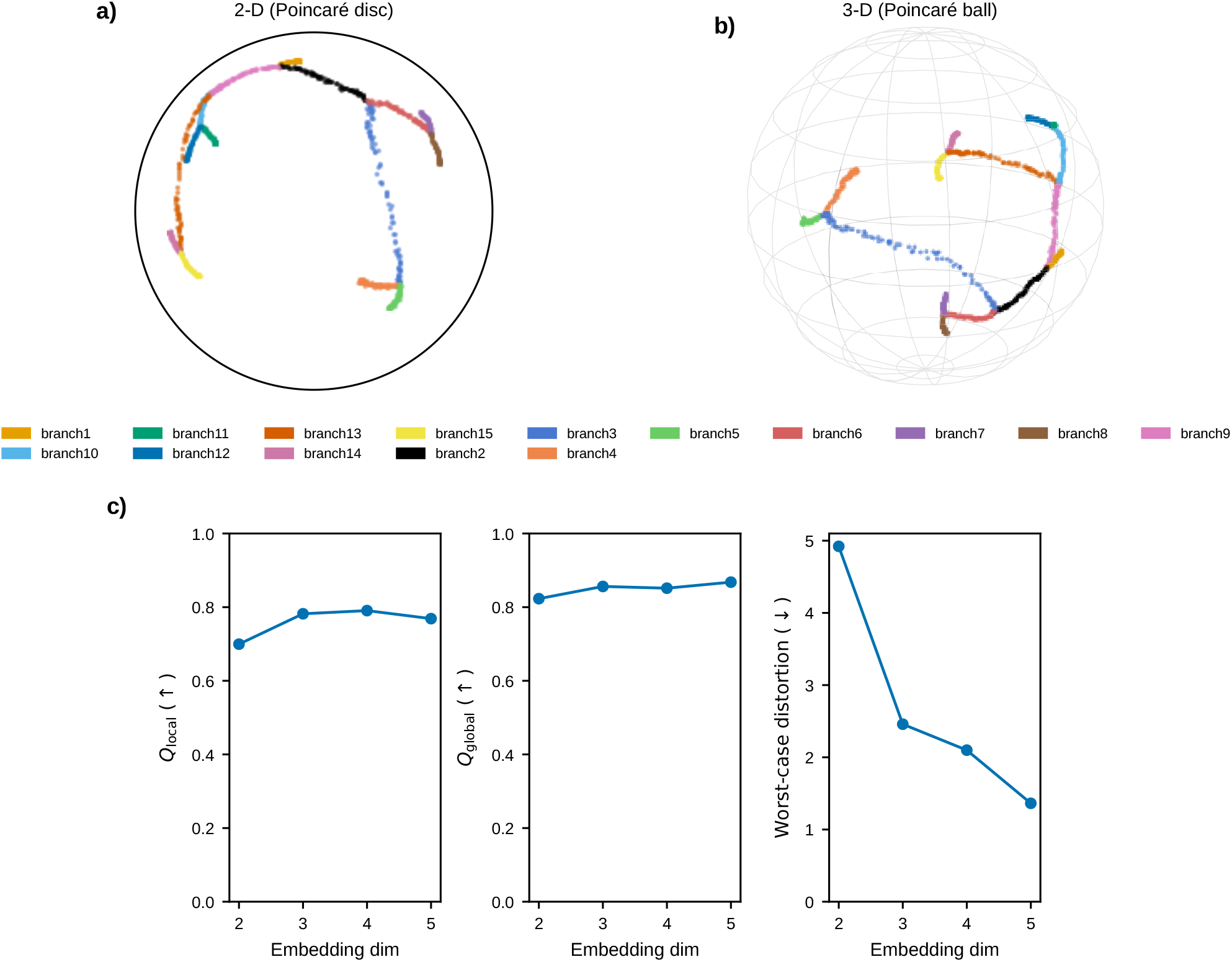
CPM beyond two dimensions. **a)** CPM embedding of the synthetic depth-3 tree in the two-dimensional Poincaré disc (*d* = 2). **b)** CPM embedding of the same data in the three-dimensional Poincaré ball (*d* = 3). **c)** Quality metrics (*Q*_*local*_, *Q*_*global*_, worst-case distortion) across the full dimensionality sweep (*d ∈ {*2, 3, 4, 5*}*), single seed, all other hyperparameters held fixed. The two-dimensional setting used throughout this study is chosen for visualization and interpretability; CPM’s architecture extends directly to higher *d* for non-visualization downstream tasks.

## B Additional results: Chicken cardiogenesis

We evaluated CPM on a multi-lineage cardiogenesis dataset (22) comprising 22,315 cells spanning myocardial, endocardial, and epicardial lineages across four developmental stages, testing whether Scanorama batch integration (23) is necessary for CPM to resolve cell-type structure. CPM embeddings colored by cell type are visually comparable with and without Scanorama batch correction (Fig. 10), and quantitatively the uncorrected embedding is at least as good: *Q*_*local*_ = 0.277, *Q*_*global*_ = 0.693, worst-case distortion = 5.24 without Scanorama, versus *Q*_*local*_ = 0.235, *Q*_*global*_ = 0.639, worst-case distortion = 8.37 with Scanorama. Batch correction therefore does not improve, and here mildly reduces, embedding quality for CPM, consistent with CPM operating directly on raw counts without preprocessing steps that can discard or distort biological signal.

**Figure 10.**
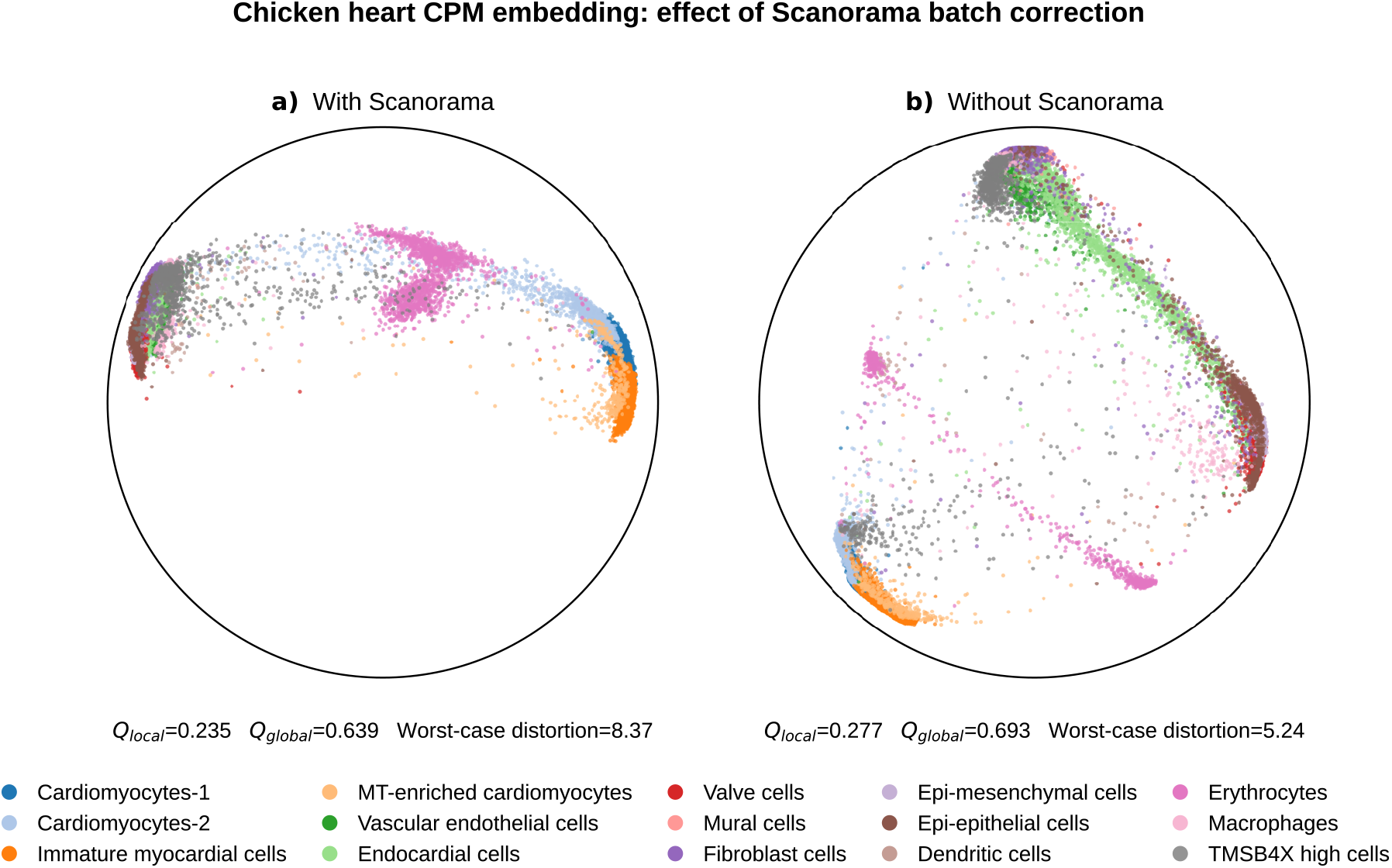
CPM on chicken cardiogenesis, with and without batch correction. **a)** CPM embedding of 22,315 cells from the chicken cardiogenesis dataset after Scanorama batch integration, colored by cell type (*Q*_*local*_ = 0.235, *Q*_*global*_ = 0.639, worst-case distortion = 8.37).**b)** CPM embedding of the same data without Scanorama batch integration, colored by cell type (*Q*_*local*_ = 0.277, *Q*_*global*_ = 0.693, worst-case distortion = 5.24).

## C Additional results: Mouse gastrulation and early organogenesis — DPT vs. CPM gene expression dynamics

We compared gene expression dynamics ordered by CPM root distance against diffusion pseudotime (DPT) along two trajectories in the mouse gastrulation atlas: definitive endoderm to gut, and visceral endoderm to gut.

On the full 116,312-cell dataset, CPM and DPT orderings along the visceral endoderm-gut trajectory both recover the expression sequence reported by Pijuan-Sala et al. (17): visceral-endoderm markers (Ttr, Apoa1) before intercalation, Pou5f1 and Sox4 during intercalation, and Hox and Cdx genes after intercalation (Fig. 11).

**Figure 11.**
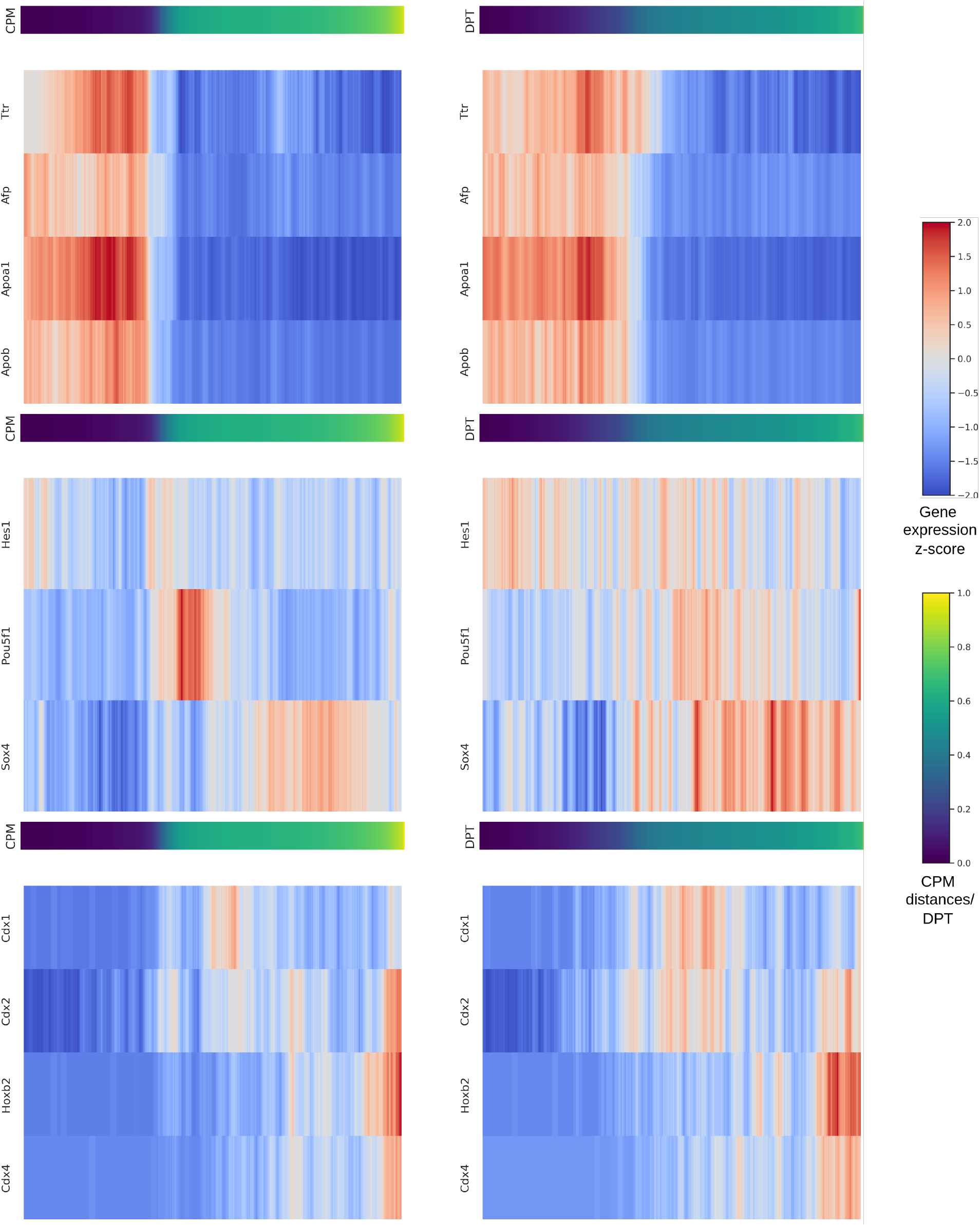
Gene expression dynamics along the visceral endoderm-gut trajectory (full dataset). CPM- and DPT-derived cell ordering reveals the initial expression of visceral-endoderm markers Ttr and Apoa1 before intercalation, followed by Pou5f1 and Sox4 dynamics during intercalation, and upregulation of Hox and Cdx genes after intercalation.

On the full dataset, DPT infers an early onset of Cdx2 and Hoxb2 along the definitive endoderm-gut trajectory; CPM root distance places their peak later, consistent with transcriptomic maturity (Fig. 12). This reflects DPT’s sensitivity to cell density: on a subsample, DPT’s visceral endoderm-gut dynamics still show persistent, artifactual Pou5f1 and Sox4 expression (Fig. 13), inconsistent with the full-dataset pattern above (Fig. 11), while its definitive endoderm-gut dynamics now align with CPM (Fig. 14). CPM’s ordering is thus more robust to cell-density variation than DPT, preserving biological resolution at full atlas scale.

**Figure 12.**
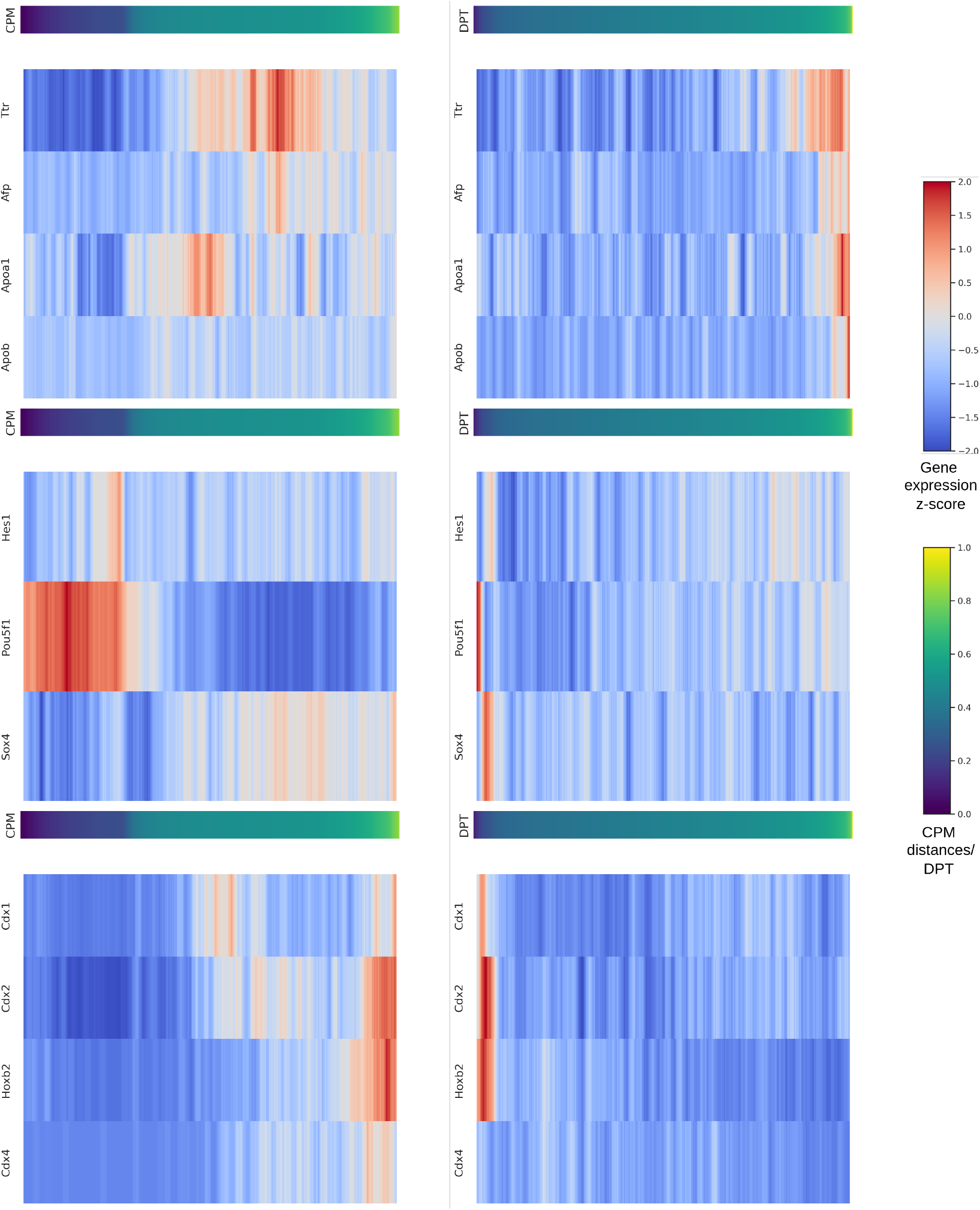
Gene expression dynamics along the definitive endoderm-gut trajectory (full dataset). DPT infers an early onset of Cdx2 and Hoxb2; ordering by CPM root distance places the peak of these factors later, consistent with transcriptomic maturity.

**Figure 13.**
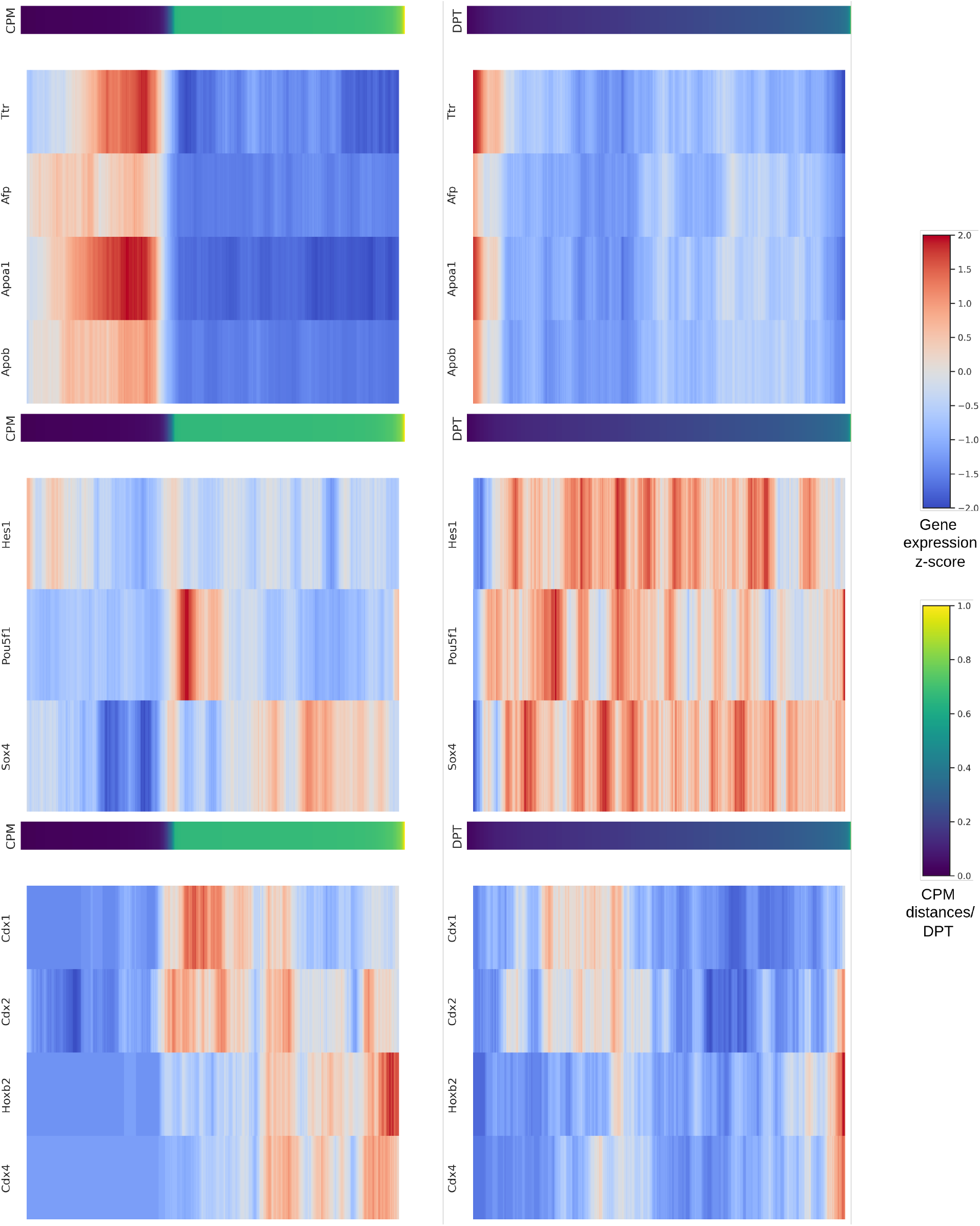
Gene expression dynamics along the visceral endoderm-gut trajectory (subsampled dataset). CPM and DPT are computed on a subsample of the dataset. DPT’s dynamics retain persistent high expression of Pou5f1 and Sox4, inconsistent with the pattern observed on the full dataset.

**Figure 14.**
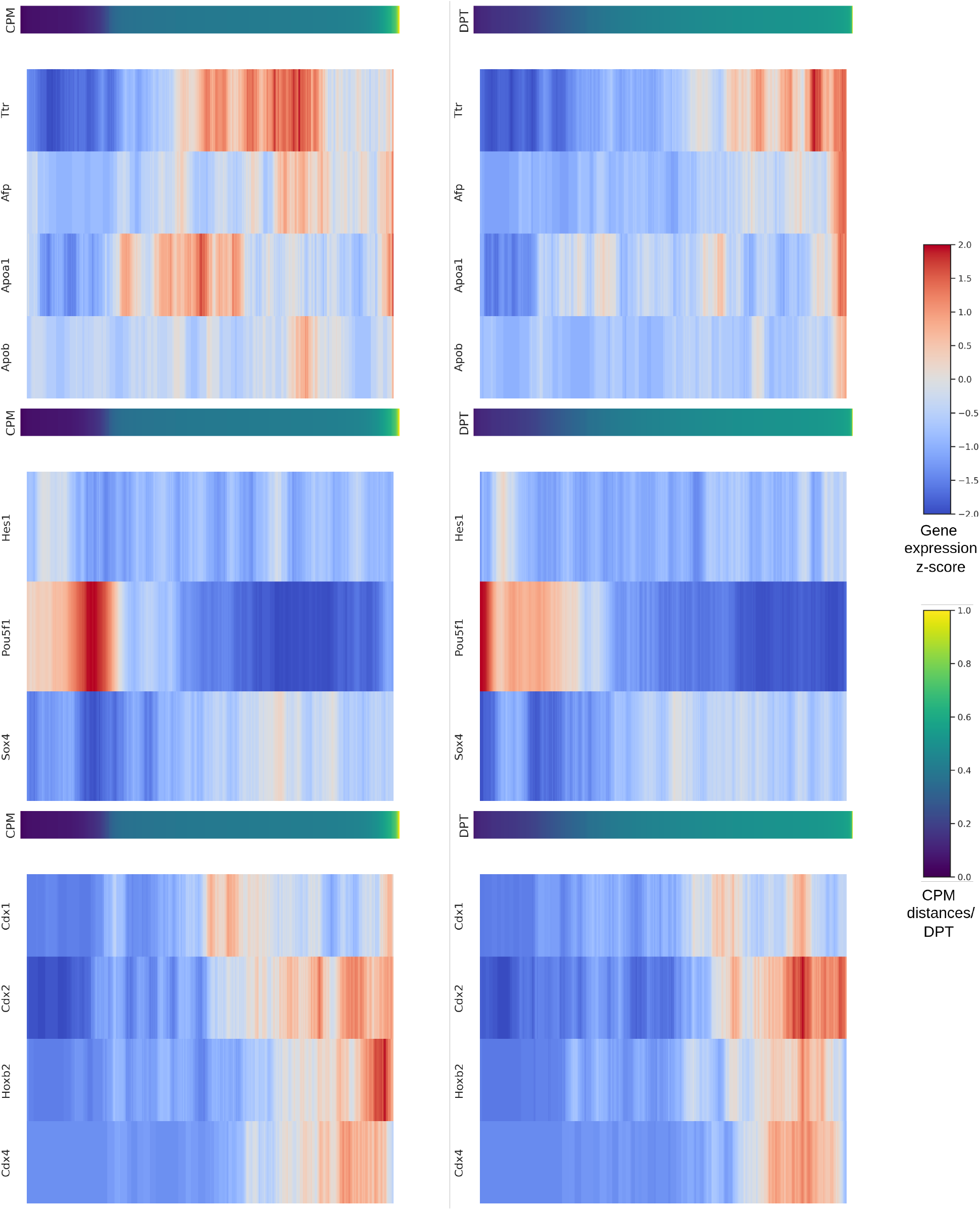
Gene expression dynamics along the definitive endoderm-gut trajectory (subsampled dataset). CPM and DPT are computed on a subsample of the dataset. DPT’s dynamics align with CPM upon subsampling and match the expected expression pattern.

## D Additional results: SOTA visualizations

This section compares CPM against the baselines on the two real developmental datasets discussed in the main text: mouse hematopoiesis (Supplementary Fig. 15) and mouse gastrulation (Supplementary Fig. 16). Because PM does not run on the full 116,312-cell mouse gastrulation dataset, we compare all baselines on a 58,156-cell subsample for this dataset instead.

**Figure 15.**
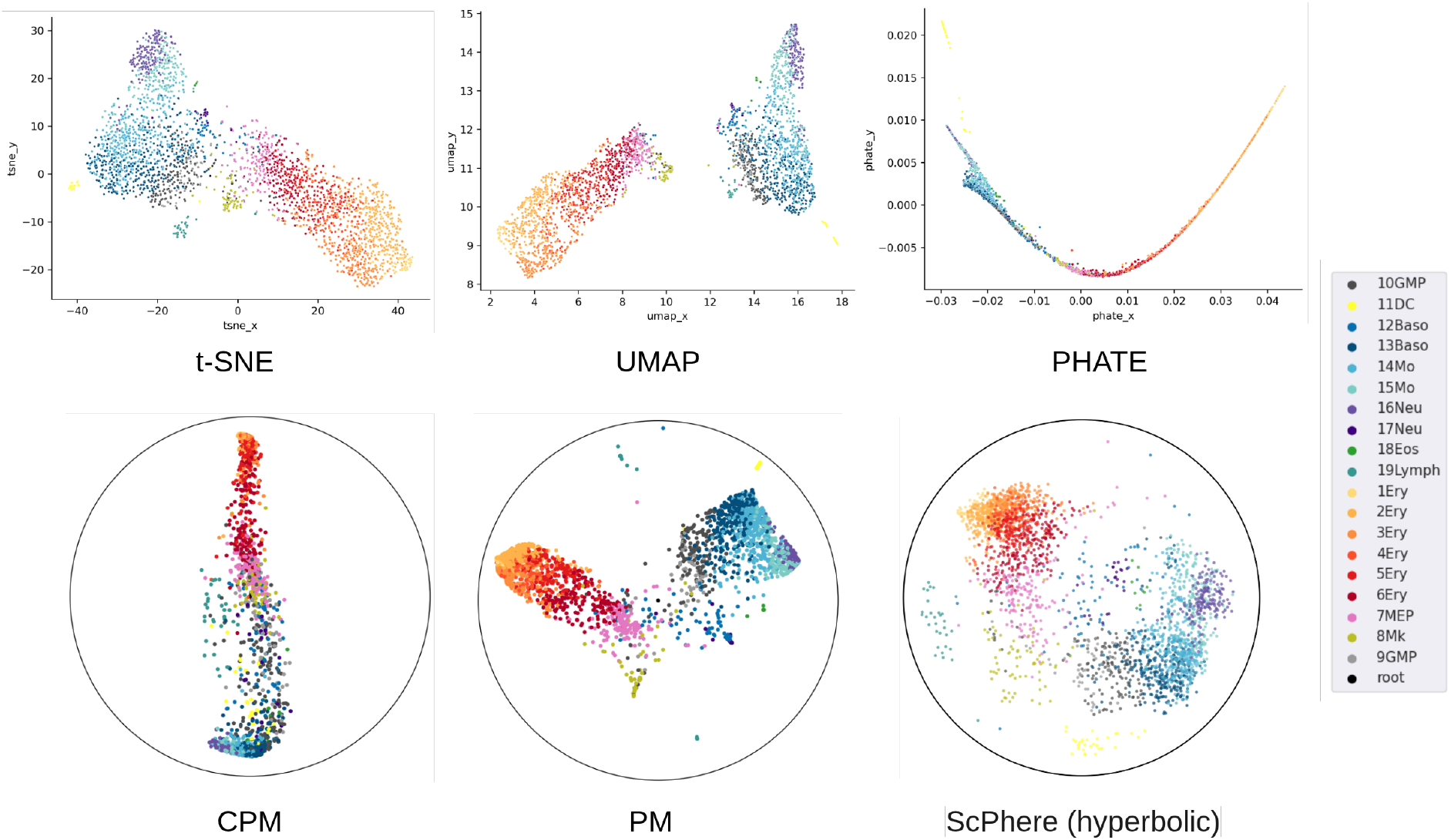
Baseline comparison on mouse hematopoiesis. Visualization of embeddings produced by the baselines on the mouse hematopoiesis dataset (2,730 myeloid progenitor cells).

**Figure 16.**
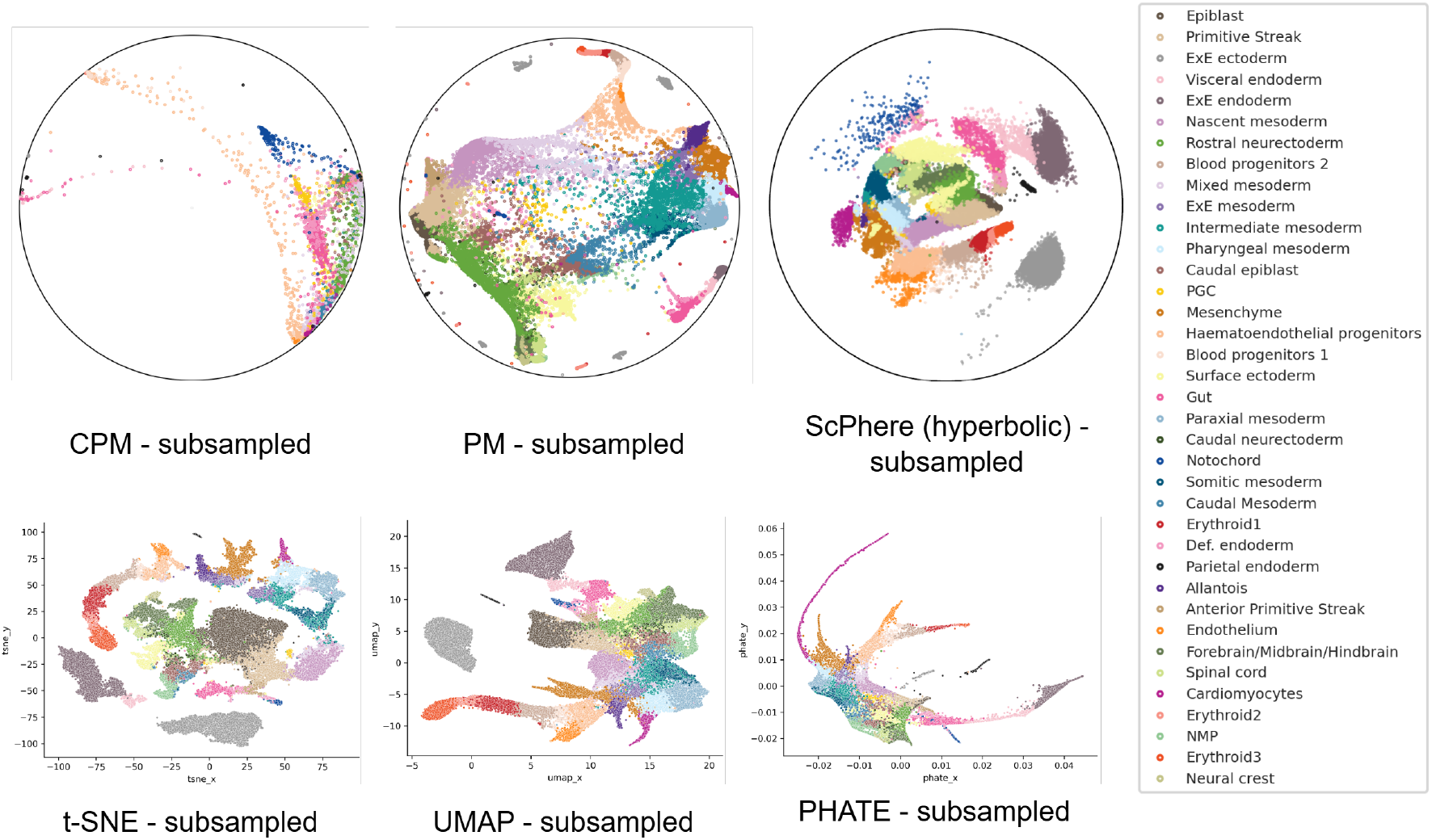
Baseline comparison on mouse gastrulation. Visualization of embeddings produced by the baselines on a 58,156-cell subsample of the mouse gastrulation dataset.

## References

(1) N. Bhasker, H. Chung, L. Boucherie, V. Kim, S. Speidel, and M. Weber. “Contrastive poincaré maps for single-cell data analysis”. In: ICLR 2024 Workshop on Machine Learning for Genomics Explorations. 2024.

(2) M. Nickel and D. Kiela. “Poincaré embeddings for learning hierarchical representations”. In: Advances in neural information processing systems 30 (2017).

(3) O. Ganea, G. Bécigneul, and T. Hofmann. “Hyperbolic neural networks”. In: Advances in neural information processing systems 31 (2018).

(4) R. Sarkar. “Low distortion delaunay embedding of trees in hyperbolic plane”. In: International symposium on graph drawing. Springer. 2011, pp. 355–366.

(5) L. v. d. Maaten and G. Hinton. “Visualizing data using t-SNE”. In: Journal of machine learning research 9. Nov (2008), pp. 2579–2605.

(6) L. McInnes, J. Healy, N. Saul, and L. Großberger. “UMAP: Uniform Manifold Approximation and Projection”. In: Journal of Open Source Software 3.29 (2018), p. 861.

(7) K. R. Moon, D. Van Dijk, Z. Wang, S. Gigante, D. B. Burkhardt, W. S. Chen, K. Yim, A. v. d. Elzen, M. J. Hirn, R. R. Coifman, et al. “Visualizing structure and transitions in high-dimensional biological data”. In: Nature biotechnology 37.12 (2019), pp. 1482–1492.

(8) A. Klimovskaia, D. Lopez-Paz, L. Bottou, and M. Nickel. “Poincaré maps for analyzing complex hierarchies in single-cell data”. In: Nature communications 11.1 (2020), p. 2966.

(9) I. Chami, Z. Ying, C. Ré, and J. Leskovec. “Hyperbolic graph convolutional neural networks”. In: Advances in neural information processing systems 32 (2019).

(10) A. v. d. Oord, Y. Li, and O. Vinyals. “Representation learning with contrastive predictive coding”. In: arXiv preprint 1807.03748 (2018).

(11) T. Chen, S. Kornblith, M. Norouzi, and G. Hinton. “A simple framework for contrastive learning of visual representations”. In: International conference on machine learning. PmLR. 2020, pp. 1597–1607.

(12) D. Bahri, H. Jiang, Y. Tay, and D. Metzler. “Scarf: Self-Supervised Contrastive Learning using Random Feature Corruption”. In: International Conference on Learning Representations. 2022.

(13) J. Ding and A. Regev. “Deep generative model embedding of single-cell RNA-Seq profiles on hyperspheres and hyperbolic spaces”. In: Nature communications 12.1 (2021), p. 2554.

(14) J. A. Lee and M. Verleysen. “Scale-independent quality criteria for dimensionality reduction”. In: Pattern Recognition Letters 31.14 (2010), pp. 2248–2257.

(15) F. Paul, Y. Arkin, A. Giladi, D. A. Jaitin, E. Kenigsberg, H. Keren-Shaul, D. Winter, D. Lara-Astiaso, M. Gury, A. Weiner, et al. “Transcriptional heterogeneity and lineage commitment in myeloid progenitors”. In: Cell 163.7 (2015), pp. 1663–1677.

(16) L. Haghverdi, M. Büttner, F. A. Wolf, F. Buettner, and F. J. Theis. “Diffusion pseudotime robustly reconstructs lineage branching”. In: Nature methods 13.10 (2016), pp. 845–848.

(17) B. Pijuan-Sala, J.A. Griiths, C. Guibentif, T. W. Hiscock, W. Jawaid, F. J. CaleroNieto, C. Mulas, X. Ibarra-Soria, R. C. Tyser, D. L. L. Ho, et al. “A single-cell molecular map of mouse gastrulation and early organogenesis”. In: Nature 566.7745 (2019), pp. 490– 495.

(18) F. A. Wolf, P. Angerer, and F. J. Theis. “SCANPY: large-scale single-cell gene expression data analysis”. In: Genome biology 19.1 (2018), p. 15.

(19) G. Bécigneul and O.-E. Ganea. “Riemannian adaptive optimization methods”. In: arXiv preprint 1810.00760 (2018).

(20) M. Weber. “Neighborhood growth determines geometric priors for relational representation learning”. In: International Conference on Artificial Intelligence and Statistics. PMLR. 2020, pp. 266–276.

(21) F. Pedregosa, G. Varoquaux, A. Gramfort, V. Michel, B. Thirion, O. Grisel, M. Blondel, P. Prettenhofer, R. Weiss, V. Dubourg, et al. “Scikit-learn: Machine learning in Python”. In: the Journal of machine Learning research 12 (2011), pp. 2825–2830.

(22) M. Mantri, G. J. Scuderi, R. Abedini-Nassab, M. F. Wang, D. McKellar, H. Shi, B. Grodner, J. T. Butcher, and I. De Vlaminck. “Spatiotemporal single-cell RNA sequencing of developing chicken hearts identifies interplay between cellular differentiation and morphogenesis”. In: Nature communications 12.1 (2021), p. 1771.

(23) B. Hie, B. Bryson, and B. Berger. “Efficient integration of heterogeneous single-cell transcriptomes using Scanorama”. In: Nature biotechnology 37.6 (2019), pp. 685–691.

